# Limited impact of the siRNA pathway on transposable element expression in *Aedes aegypti*

**DOI:** 10.1101/2024.11.26.625377

**Authors:** Alexander Bergman, Anna B. Crist, Hélène Lopez-Maestre, Hervé Blanc, Mauro Castelló-Sanjuán, Lionel Frangeul, Hugo Varet, Josquin Daron, Sarah H. Merkling, Maria-Carla Saleh, Louis Lambrechts

## Abstract

Transposable elements (TEs) are DNA sequences that can change their position within a genome. In the germline of arthropods, post-transcriptional regulation of TE expression is mainly mediated by the Piwi-interacting RNA (piRNA) pathway. piRNAs are small RNAs of 24-30 nucleotides (nt) in length produced from genomic precursor transcripts as well as through a ‘ping-pong’ amplification cycle. In somatic tissues, certain insects, such as *Drosophila*, instead rely on the small interfering RNA (siRNA) pathway as a key regulator of TE expression. siRNAs are 21nt small RNAs produced from double-stranded RNA by the endonuclease Dicer2, which guides an RNA-induced silencing complex to degrade a complementary RNA. However, whether the siRNA pathway also regulates TE expression in the mosquito *Aedes aegypti*, a medically significant vector species with abundant somatic piRNAs, is unknown. To address this question, we investigated the expression of TEs and small RNAs in both somatic and gonadal tissues of a *Dicer2* mutant line of *Ae. aegypti* and its wild-type counterpart. Our results show a modified pattern of TE expression and a decrease in TE-derived 21nt small RNAs in the *Dicer2* mutant, but no major shift of TE transcript abundance. The lack of a functional siRNA pathway also causes perturbations in piRNA ping-pong signatures and the expression of certain piRNA-associated genes, but without clear evidence for compensation by increased piRNA pathway activity. We conclude that the mosquito *Ae. aegypti* produces siRNAs targeting TEs but these lack a critical role in the regulation of TE expression both in somatic and in gonadal tissues.

## Introduction

Transposable elements (TEs), also known as transposons, are DNA sequences capable of moving within genomes (Bourque *et al*., 2018) With a few notable exceptions such as the malaria parasite *Plasmodium falciparum*, TEs are found in nearly all eukaryotic genomes in widely varying proportions (Wells and Feschotte, 2020). Among dipteran insect species, TEs have also had variable evolutionary success. The genome of the model organism *Drosophila melanogaster* is relatively poor in TEs, with a TE genome fraction of only 20% (Mérel *et al*., 2020). Among mosquitoes, the proportion of TEs in the genome is substantially higher for several species of the Culicinae subfamily, with genome fractions above 40% (Matthews *et al*., 2018; Melo and Wallau, 2020; Palatini *et al*., 2020; Ryazansky *et al*., 2024) and even over 60% for *Aedes aegypti*, compared to <20% in the Anophelinae subfamily (Compton *et al*., 2020; Melo and Wallau, 2020; Vargas-Chavez *et al*., 2022).

Depending on the presence or absence of an RNA intermediate in the transposition mechanism, TEs are divided into two classes – class I and class II. TEs that do have an RNA intermediate, such as long terminal repeat (LTR) transposons, long interspersed nuclear elements (LINEs), and short interspersed nuclear elements (SINEs) are designated class I or retrotransposons. They rely on either a self-encoded or, as is the case for SINEs, a stray reverse transcriptase (RT) to complete their transposition cycle. Class II TEs, also known as DNA transposons, such as terminal inverted repeats (TIR) transposons and helitrons, lack an RNA intermediate and transpose through a ‘cut-and-paste’ mechanism. Nonetheless, the expression of encoded proteins (e.g., transposase), allows for detection of autonomous TIR transposon expression at the RNA level, as opposed to non-autonomous TIR transposons, such as miniature inverted repeat TEs (MITEs), which only exist in DNA form (Wicker *et al*., 2007).

Due to the potentially deleterious effects of rogue transposition on genomic organization and stability, organisms have evolved various strategies to repress TE expression, such as small RNA pathways, which carry out both transcriptional and post-transcriptional silencing of TEs (Peng and Karpen, 2007; Girard and Hannon, 2008; Fagegaltier *et al*., 2009; Deniz, Frost and Branco, 2019). In *D. melanogaster,* TE expression in the germline and surrounding ovarian tissues is regulated by the Piwi-interacting RNA (piRNA) pathway. piRNAs are small RNAs of 24-30 nucleotides (nt) in length generated from genomic precursors as well as through a ‘ping-pong’ amplification loop, in which secondary piRNAs are generated through the degradation of TE transcripts (Czech and Hannon, 2016; Huang, Tóth and Aravin, 2017). In the somatic tissues, where *D. melanogaster* flies lack piRNAs, it is instead the small interfering RNA (siRNA) pathway that regulates TE expression using endogenous siRNAs (endo-siRNAs) (Chung *et al*., 2008; Czech *et al*., 2008; Ghildiyal *et al*., 2008). siRNAs are 21nt small RNAs generated from double-stranded RNA (dsRNA) by the endonuclease Dicer2, guiding an RNA-induced silencing complex (RISC) to target a complementary RNA for degradation (Zhu and Palli, 2020; Bonning and Saleh, 2021). However, observations from *D. melanogaster* do not always extrapolate to other dipteran insects. Small RNA silencing pathways have been repurposed for both somatic and germline functions throughout arthropod evolution (Lewis *et al*., 2018).

The yellow fever mosquito, *Aedes aegypti*, is an infamous vector of multiple arthropod-borne viruses (arboviruses) of medical significance, such as dengue, Zika, and chikungunya viruses (Bhatt *et al*., 2013; Aubry *et al*., 2020; Bartholomeeusen *et al*., 2023). Due to the major impact of this species on human health, the role of the siRNA pathway in *Ae. aegypti* has been previously studied in light of its antiviral function (Sánchez-Vargas *et al*., 2009; Olmo *et al*., 2018; Gestuveo *et al*., 2022; Dong and Dimopoulos, 2023; Merkling *et al*., 2023; Samuel *et al*., 2023). However, whether the siRNA pathway also regulates TEs in this species, which also has abundant somatic piRNAs (Palatini *et al*., 2017; Lewis *et al*., 2018; Joosten *et al*., 2021), remains unclear. Using a *Dicer2* (*Dcr2*) mutant line (Merkling *et al*., 2023) and an improved annotation of TEs in *Ae. aegypti* (Daron *et al*., 2024), we analyzed the transcriptomic and small RNA landscapes of the midguts and ovaries of the mutant and its wild-type control. Our findings suggest that although *Ae. aegypti* produces siRNAs targeting TEs, the siRNA pathway has an overall limited effect on TE silencing.

## Methods

### Mosquitoes

The *Dcr2*^R172fsX^ mutant line was generated as previously described (Merkling *et al*., 2023) by introducing a premature stop codon in the 5^th^ exon of the *Dcr2* gene using CRISPR/Cas9-mediated gene editing in a mosquito strain originally from Gabon. A ‘sister’ control line with the wild-type *Dcr2* gene in a shared genetic background was derived from the same crossing scheme (Merkling *et al*., 2023). Prior to the experiments, eggs were hatched in dechlorinated tap water and larvae were reared on a standard diet of Tetramin (Tetra) fish food as previously described (Merkling *et al*., 2023). After emergence, adults were maintained in insect cages (BugDorm) under a 12h-12h light-dark cycle with *ad libitum* access to a 10% sucrose solution.

### RNA extraction

Three biological replicates of 20 5-to 8-day-old females from both the *Dcr2* mutant and wild-type control lines each (16^th^ generation) were dissected into three parts – thorax, midgut, and ovaries – and snap frozen on dry ice. Total RNA was extracted from the pools with TRIzol (Life Technologies; ref. 15596026). The tissues were homogenized in 500 μl of TRIzol reagent using a Precellys homogenizer (Bertin Technologies). The RNA was phase-separated using chloroform, bound by linear acrylamide (Invitrogen; ref. AM9520) and precipitated in isopropanol. After two washes with 70% ethanol, the RNA was dissolved in 40 μl of RNase-free water. RNA concentration and purity was verified using NanoDrop (ThermoFisher Scientific). Due to excessive degradation seen in the small RNA sequencing data for thorax samples (Supplementary figure 1), these were later excluded from the analysis.

### RNA sequencing

1-2 μg of total RNA per sample were treated with DNase I (Invitrogen; AM2224), of which 200-500 ng were used for RNA-seq. RNA quality was verified using Agilent 2100 Bioanalyzer. The RNA-seq library was prepared with the Illumina Stranded mRNA Prep kit. Paired-end sequencing (2×150 cycles) was performed with a depth of 30 million paired-end reads on an Illumina NovaSeqX platform. Read quality was assessed using fastQC v0.11.9 (Andrews, 2010) and MultiQC v1.12 (Ewels *et al*., 2016).

### RNA-seq data analysis

RNA-seq data were analyzed using the rnaflow pipeline (Legendre, 2024). In brief, raw reads were trimmed using cutadapt v2.10 (Martin, 2011) using the parameters ‘-m 25 –O 6 ––trim-n ––max-n 1 –q 30’. Reads were aligned to the AaegL5 *Ae. aegypti* reference genome (VectorBase release 61) (Alvarez-Jarreta *et al*., 2024) using STAR v2.7.9a (Dobin *et al*., 2013) two-pass alignment with the options ‘––outFilterMismatchNoverLmax 0.05 ––outFilterMultimapNmax 50’. Reads were counted using TEtranscripts v2.2.3 (Jin *et al*., 2015) using ‘––mode multi’. A principal component analysis (PCA) of variance-stabilized counts was done using the R package FactoMineR v2.9 (Lê, Josse and Husson, 2008). Differential expression analysis was performed on a concatenated table of gene and TE counts using DESeq2 v1.40.2 (Love, Huber and Anders, 2014) for each tissue separately. Detection of chimeric reads was done using ChimeraTE v1.1.1 (Oliveira *et al*., 2023). For transcripts per million (TPM) values, read counts were normalized by the length of collapsed exons for genes and the family consensus length for TEs. Gene set enrichment analysis was performed on the DESeq2 output using the R package fgsea v1.26.0 (Korotkevich *et al*., 2021) on genes and TEs separately. For TEs, individual families were grouped by their order. Genes were grouped according to KEGG (Kanehisa and Goto, 2000) (release 106) pathway annotations. Three additional gene sets were added, containing either siRNA-, piRNA-, or histone modification-related genes based on homology to selected *D. melanogaster* gene ontology (GO) terms. Homology was determined using annotation from VectorBase release 66. Ensembl gene IDs were transformed into *AAEL*-nomenclature using the online tool DAVID (Sherman *et al*., 2022). All plots were made using the R package ggplot2 v3.4.4 (Wickham, 2016). The threshold for statistical significance for this and all other analyses was set to p <0.05.

### Small RNA sequencing

5 μg of total RNA per sample were used directly for small RNA sequencing. Small RNAs of 19-33 nt in length were purified from 15% acrylamide/bisacrylamide (37.5:1), 7 M urea gel as previously described (Gausson and Saleh, 2011). The small RNA library was prepared using NEB Next Multiplex Small RNA Library Prep kit for Illumina (New England Biolabs [NEB]; Ipswitch, MA, USA; ref. E7300 L) with Universal miRNA Cloning Linker (NEB; ref ES1315S) as the 3’ adaptor and in-house designed indexed primers. Libraries were diluted to 4 nM and sequenced using an Illumina NextSeq500 High Output kit v2 (75 cycles) on an Illumina NextSeq500 platform over 52 cycles.

### Small RNA-seq data analysis

Raw reads were pre-processed through poly-A/T/C/G removal, adapter trimming and filtering for contaminating long RNA using cutadapt v2.10 (Martin, 2011) with options ‘-e 0.15 –O 6 ––trimmed-only –m 18 ––match-read-wildcards –q 20’.

For 21nt RNA analysis, pre-processed reads were aligned to the AaegL5 *Ae. aegypti* reference genome (VectorBase release 61) using bowtie v1.2.3 (Langmead *et al*., 2009) using the options ‘-v 1 –a –M 1’ (random attribution of multimapping reads). Reads that mapped to annotated micro-RNAs (miRNAs), transfer RNAs (tRNAs), small nuclear RNAs (snRNAs), and small nucleolar RNAs (snoRNAs), as well as an unannotated miRNAs, aae-miR-989 (Li *et al*., 2009) and a gene (AAEL019428) containing a highly expressed aae-miR-2942 precursor, previously detected (Su *et al*., 2017) in *Ae. albopictus* as aal-miR-956p with one mismatch, were filtered out. To filter out 21nt piRNA fragments from the alignment, pre-processed reads were collapsed and dusted using the small RNA NGS toolbox v2.1 (Rosenkranz *et al*., 2015), mapped to the reference genome using sRNAmapper v1.0.5 (Rosenkranz *et al*., 2015) (option ‘-alignments best’) and used to annotate piRNA clusters using proTRAC v2.3.1 (Rosenkranz and Zischler, 2012) with default options. piRNA clusters were annotated for each sample and combined for each tissue and condition (*Dcr2* mutant or wild-type) pair. 21nt reads mapping to annotated clusters were filtered out from the alignment. Finally, 21nt RNAs were counted using TEtranscripts v2.2.3 (option ‘––mode multi’) (Jin *et al*., 2015). Reads per million (RPM) values for each order of TEs were normalized by the total miRNA RPM, under the assumption that the size of the total miRNA pool remains constant within a given tissue across replicates and conditions. For TPM values, the read counts were normalized against the length of collapsed exons for genes and the family consensus length for TEs and subsequently normalized by the total miRNA TPM.

### Ping-pong signature analysis

The size range of potential piRNAs was defined to 26-30 nt based on read size distribution (Supplementary figure 2). For piRNA analysis, pre-processed reads were aligned using bowtie v1.2.3 (Langmead *et al*., 2009) using the options ‘-v 1 –a ––best –strata’ and filtered for annotated small RNA genes. The alignments were then subset by TE order and strand with relation to the annotated TE copy and filtered for reads mapping to more than one orientation relative to all copies within the order or more than one order of TEs. Reads mapping to the antisense strand were considered potential primary piRNAs and intersected with reads mapping to the sense strand, considered potential secondary piRNAs. Overlapping pairs where the potential primary piRNA was upstream of the potential secondary piRNA on the sense strand of the TE were counted and normalized by the number of mappings of each member of the pair. Logo plots of potential and putative ping-pong (i.e., overlapping by 10 nt) primary and secondary piRNAs were generated using the plot_seqlogo functionality of the biopieces package (Hansen, unpublished). Counts for sense, antisense, and putative primary and secondary reads were generated by extracting unique read names from the subset alignments. Further investigation of putative secondary piRNAs was conducted by separating secondary piRNA reads based on which set of copies they map to in order to identify the sources of ping-pong signatures.

### Reverse transcriptase activity assay

RT activity was measured in midguts and ovaries using a protocol adapted from Wu *et al*., 2021 and Pyra, Böni and Schüpbach, 1994. Four pools of 5 tissues from the *Dcr2* mutant line and the wild-type control line (24^th^ generation) were collected at 5-7 days post adult emergence, with midguts and ovaries originating from the same mosquitoes. Organs were dissected and placed in 100 μL of CHAPS lysis buffer (Goic *et al*., 2013), containing 10 mM Tris-HCl pH 7.5 (Invitrogen; ref. AM9855G), 400 mM NaCl (Invitrogen, ref. AM9759), 1 mM MgCl_2_ (Invitrogen; ref. AM9530G), 1 mM EGTA (Sigma-Aldrich; ref. E4378), 0.5% CHAPS (ThermoFisher Scientific; ref. 28300), 10% glycerol (Sigma-Aldrich; ref. J61059-AP), 1 mM DTT (Invitrogen; ref. Y00147), and 1X cOmplete EDTA-free protease inhibitor cocktail (PIC) (Roche; ref. 11873580001). The tissues were homogenized in a Precellys homogenizer (Bertin technologies) and stored at –80°C. Subsequently, the samples were clarified by centrifugation for 10 minutes at 16,000*g* to remove tissue debris. The resulting supernatant was collected into a new tube. Protein content in the supernatant was measured using Pierce Detergent Compatible Bradford Assay kit (ThermoFisher Scientific; ref. 23246) according to the manufacturer’s instructions. All samples were diluted to the protein concentration of the least concentrated sample. Prior to the RT reaction, 30 pmol of MS2 reverse primer (5’CGCTTGTAGGCACCTTGATC) with 0.4 mM dNTPs (ThermoFisher Scientific; ref. R1121) were allowed to anneal to 100 ng of MS2 RNA (Roche; ref. 10165948001) in a 12.5-μL annealing reaction at 65°C for 5 minutes. The annealing reactions were used in subsequent 25-μL RT reactions, containing 30 pmol MS2 reverse primer, 100 ng MS2 RNA, 0.2 mM dNTPs, 1 mM DTT (Invitrogen; ref. Y00147), 10 mM Tris-HCl pH 7.5 (Invitrogen; ref. AM9855G), 1 mM KCl (Invitrogen; ref. AM9640G), 0.14 mM MnCl_2_ (Sigma-Aldrich; ref. M1787), 0.02% Triton X-100 (Sigma-Aldrich; ref. 93433), 40 units of RNase OUT (Invitrogen; ref. 10777-019) and 5 μL of protein sample. The RT reaction was allowed to progress for 1 hour at 25 °C, after which it was stopped by heat-inactivation at 70°C for 15 minutes. Heat-inactivated samples (45 minutes of incubation at 98°C), CHAPS buffer alone, and no-template reactions served as negative controls. Three technical replicates of 1 unit of SuperScript II (Invitrogen; ref. 18064022) were used for relative quantification of RT activity. Generated MS2 cDNA was quantified with two technical replicates through quantitative PCR (qPCR) in a 10-μL reaction containing 3 pmol MS2 forward primer (5’TCCCGGTCGTTTTAACTCGA), 3 pmol MS2 reverse primer (5’CGCTTGTAGGCACCTTGATC), and 1 μL of RT reaction (template) using Promega GoTaq qPCR Master Mix (Promega; ref. A600A). Statistical analysis was done using a linear mixed effect model using the R packages lme4 v1.1-35.1 (Bates *et al*., 2015) and lmerTest v3.1-3 (Kuznetsova, Brockhoff and Christensen, 2017) with organ and condition as explanatory variables and biological replicate within each condition as a blocking variable (degrees of freedom estimation method: Kenward-Roger). P-values for pairwise comparisons were calculated using Wald’s test through the package emmeans v1.9.0 (Lenth, Russel V., 2023).

## Results

To determine the role of the siRNA pathway in TE expression regulation in *Ae. aegypti*, we analyzed the abundance of TE transcripts and small RNAs in the midgut and the ovaries of the *Dcr2* mutant and its wild-type control. We excluded non-autonomous and unknown TEs from the main analysis due to frequent chimerism with gene transcripts (Supplementary figure 3) and limited significance. Nonetheless, we provide the full results in Supplementary figures 4, 5, 6, 7, 9, 10, 12, and 13.

### *Dcr2* mutant displays specific differences but no uniform shift in TE expression

We first compared the abundance of gene– and TE-derived transcripts between the *Dcr2* mutant and the wild-type control. In total, the RNA-seq libraries yielded between 15.1 million and 30.6 million read counts per biological replicate. Of these, approximately 1% in ovary samples and 2% in midgut samples originated from TEs (Figure 1A). The proportion of TE counts was significantly influenced by the organ but not by the mosquito line (ANOVA on logit-transformed proportions, p = 0.79 for line, p = 6.4×10^-8^ for organ, non-significant interaction excluded from the model). A PCA confirmed the clear distinction between the tissues in terms of both gene and TE expression, with the first component explaining 82% and 75% of the variation, respectively (Figure 1B). However, the PCA also showed a clear separation of the two conditions along the second axis, which explained 4.3% of the variation for genes and 8.3% of the variation for TEs. Differential expression analysis revealed that specific TE families were both enriched and depleted between the lines for every order of autonomous TEs (Figure 2A). In the midgut, 24 families were enriched and 60 families depleted with an absolute log_2_ fold change greater than 1 at a significance level of adjusted p <0.05. Under the same criteria, 82 families were enriched and 57 depleted in the ovaries. Nonetheless, gene set enrichment analysis (GSEA) found that no TE order was differentially expressed as a whole (Figure 2B). Functional validation using an RT activity assay followed the trends seen in the GSEA analysis for LINE and LTR transposons, with a trend toward lower RT activity in the midgut and higher RT activity in the ovaries (Figure 2C). The linear mixed effect model showed a significant interaction between organ and line (p = 0.008), reflecting the opposite effects of the *Dcr2* mutation on RT activity in the midgut and the ovary. Together, these data provide evidence that differences in the expression of specific TE families do exist in the absence of *Dcr2*, but they are not consistent across orders or tissues in their direction or their magnitude.

**Figure 1.**
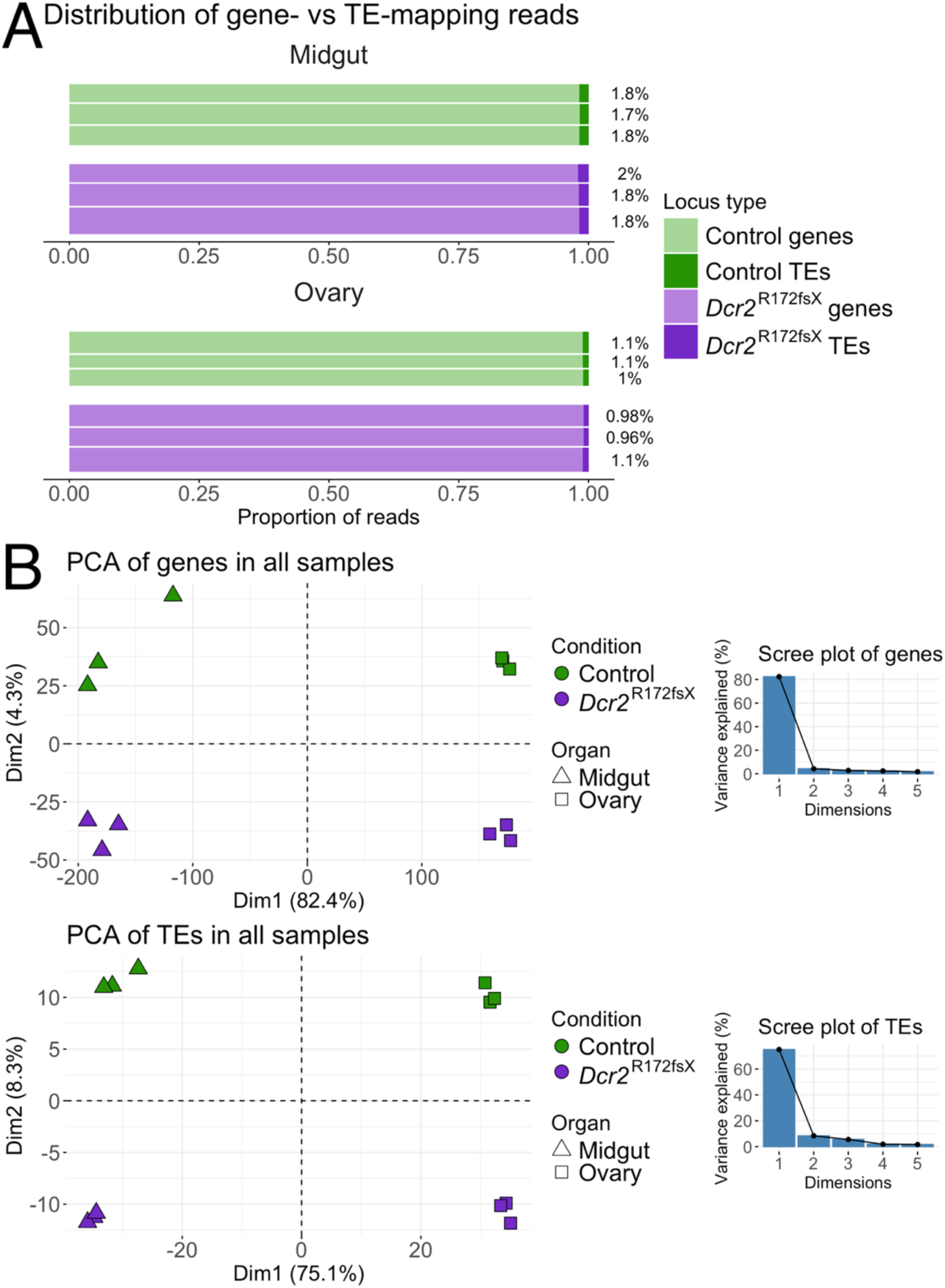
– TE and gene expression patterns differ in the *Dcr2* mutant. (A) Proportion of reads mapping to either genes or TEs in the RNA-seq libraries from midguts (top graph) and ovaries (bottom graph). The vertical height of the bar is proportional to library size (number of counted reads). Percentages of TE-mapping reads are stated on the right side. (B) Separate principal component analyses (PCAs) of gene (top graph) and TE (bottom graph) read counts with accompanying scree plots (line plots of the eigenvalues of principal components).

**Figure 2.**
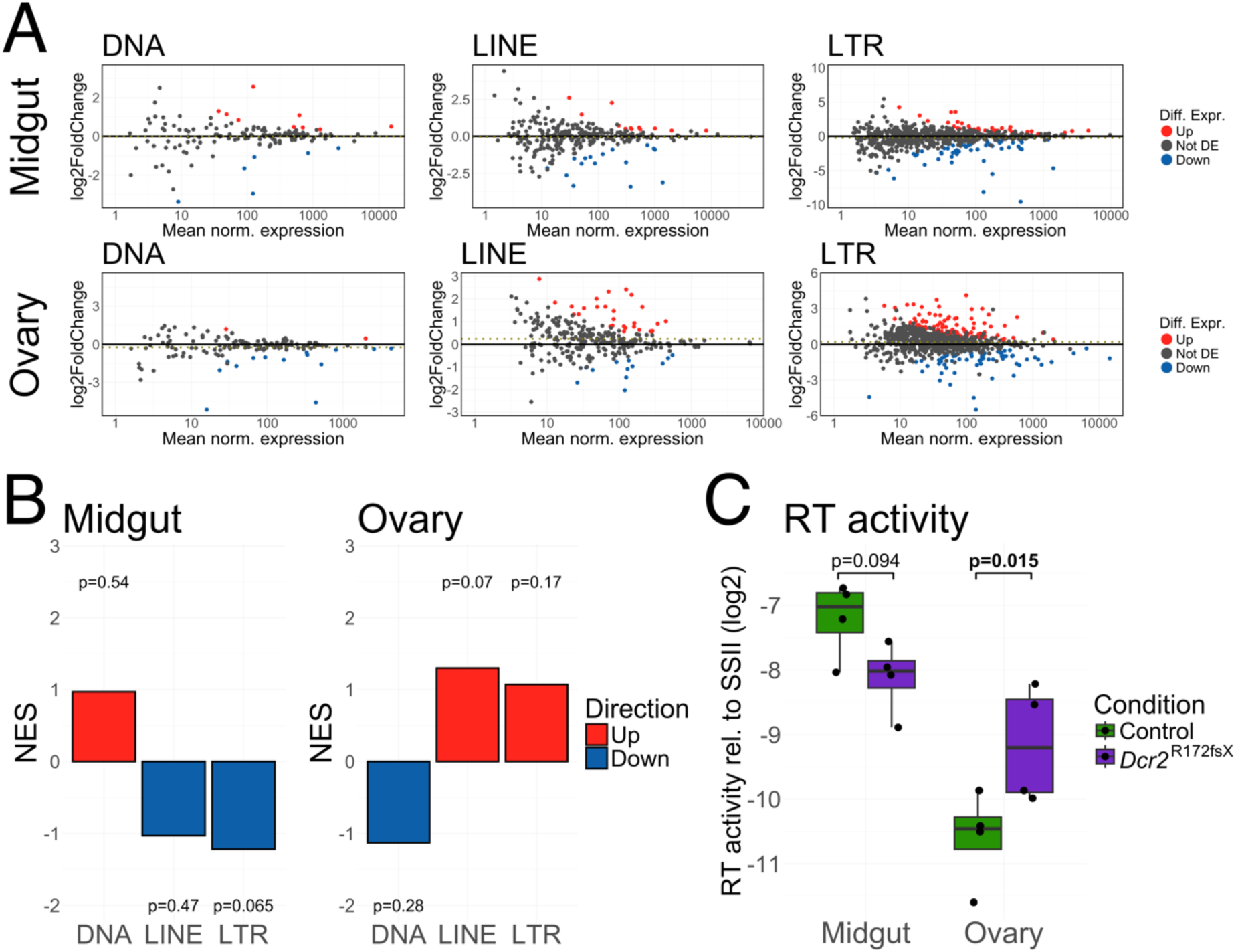
– TE expression in the *Dcr2* mutant is perturbed but not uniformly shifted. (A) MA plots of individual TE families grouped by order (DNA, LINE, LTR) for midguts (top row) and ovaries (bottom row). The x-axis shows mean read counts normalized by the median of ratios (DESeq2-based normalization) and the y-axis shows log_2_ fold-change in the *Dcr2* mutant line. Families are colored according to their differential expression (red: enriched in mutant line; blue: depleted in mutant line; grey: not differentially expressed). The dotted line in the center of each plot represents the mean log_2_ fold-change. (B) Gene set enrichment analysis results. The height of each bar represents the normalized enrichment score (NES), i.e., the relative enrichment of the TE order compared to a random, equally sized, group of genes. P-values above or below the bars indicate the false discovery rate for the enrichment (red bars) or depletion (blue bars) in the *Dcr2* mutant relative to the wild-type control. (C) Box plot of RT activity measured in midgut and ovary samples relative to SuperScript II (SSII). P-values shown above the graph were generated by pairwise comparisons within a linear mixed effect model using Wald’s test.

### Reduction of TE-derived 21nt RNAs does not correlate with expression differences

To determine whether the lack of a major shift in TE transcript abundance in the absence of *Dcr2* was simply due to a general lack of TE targeting by the siRNA pathway, we examined the abundance of TE-derived 21nt small RNAs. We mapped small RNAs to the genome with random attribution of multimapping reads to retain the correct total number of reads. Misattribution through randomization was rare, with at most 1.5% and 3.1% of multimapping 21nt reads mapping to more than one TE family in midgut and ovary samples, respectively. We scaled RPM values by total miRNA RPM to adjust for the change in proportion following a change in composition of small RNA-seq data, following the assumption of a constant size of the miRNA pool. In the *Dcr2* mutant, all autonomous TE orders exhibited a significant reduction in the abundance of TE-mapping 21nt RNAs except LTR transposons in the ovaries, for which the difference was marginally non-significant (Figure 3A). This result confirms that TE transcripts are collectively targeted by the siRNA pathway. To explain the discrepancy between the overall depletion of siRNAs and the lack of a major shift in TE transcript abundance in the *Dcr2* mutant, we hypothesized that only some TE families within each order were disproportionately affected by a dysfunctional siRNA pathway. Under this hypothesis, we predicted that the TE families with the greatest abundance of 21nt RNAs (i.e., the most targeted TE families) in the wild-type line would display the highest enrichment in the *Dcr2* mutant. The analysis of ratios between 21nt RNA and transcript abundances by TE family did not support this hypothesis. Instead, we found that TE families whose expression was depleted in the *Dcr2* mutant line tended to display the lowest amounts of TE-derived 21nt RNAs in the wild-type line (Figure 3B). The families enriched in the *Dcr2* mutant were thus not those subject to the most intense targeting in the wild-type control. Overall, we found that in both the midgut and the ovaries, fewer TE-derived 21nt RNAs were detected in the *Dcr2* mutant but the reduction did not correlate with differential TE expression.

**Figure 3.**
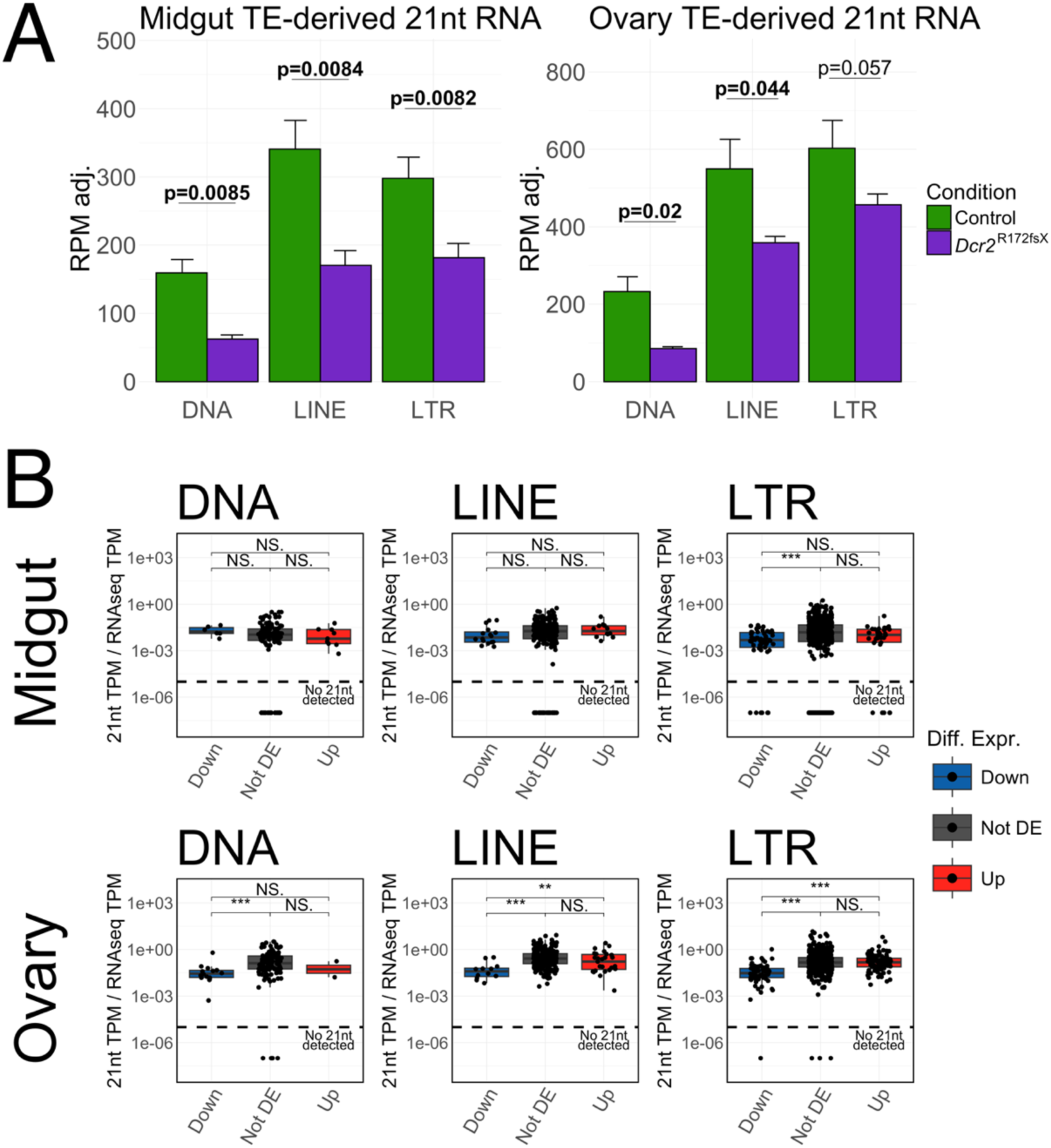
– Reduction of TE-derived 21nt RNAs in the *Dcr2* mutant does not correlate with differential TE expression. (A) miRNA-adjusted reads per million (RPM) mapping to the different TE orders in the midgut (left plot) and ovaries (right plot) of *Dcr2* mutant and control mosquitoes. Error bars denote one standard deviation. P-values shown above the bars were generated with Welch’s t-test. (B) Ratios between miRNA-adjusted 21nt RNAs expressed in transcripts per million (TPM) and RNA-seq TPM in the control mosquitoes for individual TE families depleted (Down), non-differentially expressed (Not DE), and enriched (Up) in the *Dcr2* mutant within each TE order for midguts (top row) and ovaries (bottom row). P-values shown above the graphs were generated using Wilcoxon rank sum test (*<0.05, **<0.01, ***<0.001, NS = non-significant). Families with no detected transcripts were excluded. Families with detected transcripts but no detected 21nt RNAs are shown below the dashed line.

### Lack of *Dcr2* only causes a minor change in piRNA ping-pong activity in the midgut

Since a reduction in TE-mapping 21nt RNAs was not associated with a corresponding increase in transcript abundance, we investigated whether there was any compensation through increased ping-pong activity of the piRNA pathway. In the ovary, canonical signatures of ping-pong amplification could be observed for the three autonomous TE orders (DNA, LINE, and LTR) in both control and mutant mosquitoes, with both frequent 10nt overlaps and a corresponding 1U and 10A bias in putative primary and secondary piRNAs, respectively (Figure 4). In the midgut, evidence of ping-pong activity, consisting of frequent 10nt overlaps and 1U/10A biases could only be seen for DNA and LTR transposons (Figure 4). For DNA transposons, the overabundance of 10nt overlaps varied between biological replicates (Figure 4). Furthermore, the number of putative secondary piRNAs was negligible (8-167 reads) and specific sequences dominated putative primary piRNAs involved in ping-pong amplification (Supplementary figure 4). The importance of the ping-pong cycle in DNA transposon regulation is therefore likely minimal. The only significant difference that we observed between the two mosquito lines was a decrease in 10nt overlaps for LTR transposons in the midgut of the *Dcr2* mutant (Figure 4). However, a 10A and 1U bias was seen for LTR-derived putative piRNAs in both conditions (Figure 4), and no differences were observed in the amounts of putative secondary (Supplementary figure 5) nor primary (Supplementary figure 6) piRNAs. Closer inspection revealed that the vast majority of the piRNA pairs overlapping by 10nt in the wild-type mosquitoes originated from a single TE copy belonging to the family TE_0669_Gypsy (Supplementary figure 7), the secondary piRNA-component of which was mainly composed of a single sequence (Supplementary figure 8). Although 10nt overlaps from this family are also abundant in the *Dcr2* mutant, these originate from multiple other TE copies and do not account for as many of the 10nt-overlapping pairs (Supplementary figure 7). Removing this single locus from the analysis abrogated any difference between the two conditions (Supplementary figure 9). Furthermore, this analysis showed that the ping-pong activity for LTR transposons in the midgut was largely driven by a handful of Gypsy superfamily members (Supplementary figure 7A), with ping-pong activity being low in the midgut in general (Supplementary figures 4, 5, 6). Overall, we only detected a change in ping-pong signatures for LTR transposons in the midgut but this difference was attributable to a single locus. We found no generalized compensation by the piRNA pathway for the lack of *Dcr2*.

**Figure 4.**
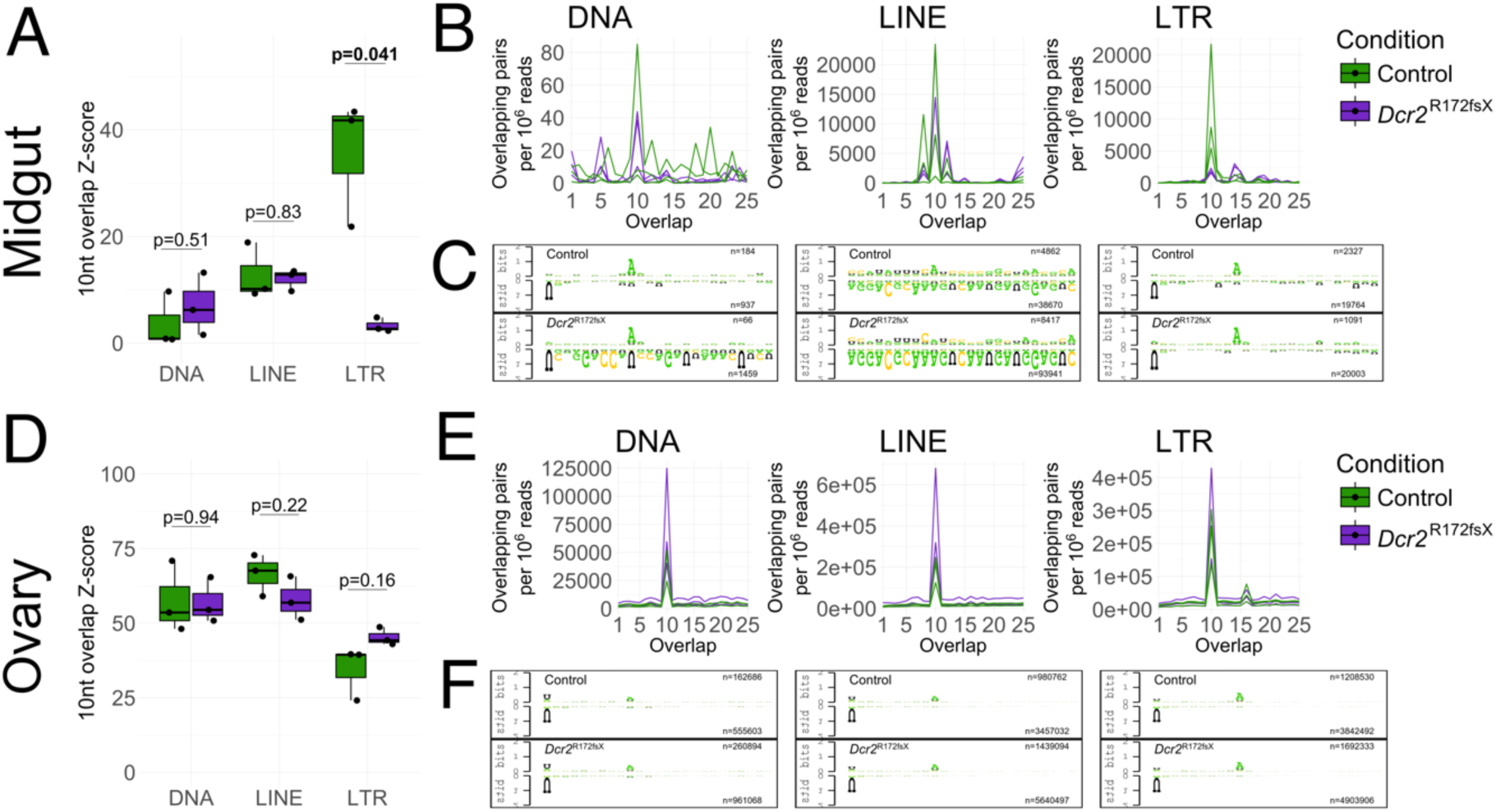
– TE-derived piRNAs display mostly unchanged ping-pong activity in the *Dcr2* mutant. (A, D) Box plots of 10nt overlap Z-scores among 26-30nt sense and antisense reads mapping to TEs stratified by order in midguts (A) and ovaries (D) of the *Dcr2* mutant and the wild-type control. P-values indicated above the box pairs were generated using Welch’s t-test. Significant p-values are highlighted in bold font. (B, E) Frequency of overlaps among sense and antisense 26-30nt reads by a given number of nt for midguts (B) and ovaries (E). (C, F) Logo plots of sense (“secondary”, top logo) and antisense (“primary”, bottom logo) 26-30nt reads overlapping each other by 10nt for control (top row) and *Dcr2* mutant (bottom row) samples for midgut (C) and ovary (F). The sequences from all three replicates were merged into one logo and trimmed to 25nt. The number of sequences used to construct the logos is specified on the right side within each plot. For complete and replicate-specific logo plots, see supplementary figure 4.

### Transcriptomic changes show no obvious compensatory mechanism in *Dcr2* mutant

To investigate whether the lack of *Dcr2* was associated with another compensatory mechanism than piRNA ping-pong activity that could contribute to TE expression regulation, we performed a differential gene expression analysis of the RNA-seq data. In total, 413 genes (127 enriched, 286 depleted) in the midgut and 1234 genes (546 enriched, 688 depleted) in the ovaries were differentially expressed in the *Dcr2* mutant with an absolute log_2_ fold-change greater than 1 below the significance threshold of an adjusted p <0.05 (Supplementary figure 10). In the ovaries, several metabolic pathways were downregulated, while pathways related to nuclear processes, such as transcription factors and spliceosome-associated genes, were enriched. In the midgut, the only significantly differentially regulated pathway was nucleotide excision repair (Supplementary figure 11A). A gene-by-gene analysis of the differential expression of siRNA-, piRNA-, and histone modification-related genes showed a depletion of certain factors involved in piRNA biogenesis, such as *Yb*, *papi*, and *vas* homologs, as well as *Piwi1/3* and *Piwi2* in the ovaries. Interestingly, expression of *Dicer1*, canonically an essential gene for miRNA biogenesis, was slightly elevated (Supplementary figure 11B). Overall, the *Dcr2* mutation caused significant perturbation in the transcriptome homeostasis of the ovaries, but no single compensatory mechanism could be discerned from this dataset.

## Discussion

In the present study, we compared the transcriptomic and small RNA landscapes between wild-type and *Dcr2*-deficient *Ae. aegypti* and showed that the lack of a functional *Dcr2* allele has wide and multifaceted effects on gene and TE expression in both midguts and ovaries in this genetic background. Our results confirm previously published detection of TE-derived endo-siRNAs in *Ae. aegypti* (Arensburger *et al*., 2011; Ma *et al*., 2021), but their function as regulators of TE transcript abundance had not been studied until now. Despite a reduction of putative siRNAs originating from TEs in the *Dcr2* mutant (Figure 3A), we did not observe a major enrichment of transcripts from autonomous TEs. Our data challenge the assumption that the siRNA pathway is a key regulator of TE expression in *Ae. aegypti*.

In *D. melanogaster*, a disrupted siRNA pathway is associated with an increase in TE expression in somatic tissues, even though not all TE families may be affected equally (Chung *et al*., 2008; Czech *et al*., 2008; Ghildiyal *et al*., 2008; Mirkovic-Hösle and Förstemann, 2014; Roy *et al*., 2020). In this study, somatic expression of TEs was not affected, and, if anything, trending downward for retrotransposons. Our data show numerous significant changes in TE transcript abundance in both directions, suggesting highly varying effects of a dysfunctional siRNA pathway on the expression of specific TE families (Figure 2A; Supplementary figure 10). However, combining the RNA-seq and small RNA-seq datasets show that the TE families that are more expressed in the *Dcr2* mutant are not more intensely targeted by the siRNA pathway in the wild-type control (Figure 3B). This either suggests that certain TE families are more sensitive to siRNA-mediated silencing and thus become enriched in the *Dcr2* mutant despite levels of siRNA targeting in the wild-type control being akin to those of less sensitive families, or that the enrichment of these families is due to an indirect effect of the *Dcr2* mutation. Beyond its role in RNA interference, Dicer2 has been shown to have a role in immune gene expression and cytoplasmic polyadenylation in insects (Wang *et al*., 2015; Coll *et al*., 2018; Dong *et al*., 2022). Furthermore, abundance of transcript is not necessarily equivalent to autonomous expression, as TEs are commonly found in chimeric transcripts and could be affected by the expression of nearby genes, just as they themselves can act as *cis*-regulators (Kapusta *et al*., 2013; Chuong, Elde and Feschotte, 2017; Treiber and Waddell, 2020). Elucidating whether differential expression of TE families arises from differential sensitivity to siRNA-mediated silencing, a side effect of the *Dcr2* mutation, or a by-product of differential gene expression would require further studies on the reliance of individual TE families on specific TE regulation mechanisms in *Ae. aegypti*.

We did not find evidence of compensation for the lack of siRNAs through increased piRNA ping-pong activity. The only difference we observed between the *Dcr2* mutant and the control mosquitoes across the piRNA pathway was in the midgut, but (i) in the opposite direction of hypothetical compensation and (ii) originating from a single locus (Figure 4; Supplementary figures 5, 6, 7, 9). In general, although piRNAs are readily detectable in the midgut of *Ae. aegypti*, ping-pong activity appears restricted to a handful of TE families.

In the ovaries, we found that the *Dcr2* mutation had an effect opposite to the trend in the midgut, with increased RT activity and an upward trend in retrotransposon transcription. Despite that, we did not observe a change in ping-pong signature other than a weak downward trend in piRNA abundance (Supplementary figures 5, 6). We noticed that several genes important for piRNA biogenesis were depleted, although the piRNA-related gene set as a whole was not significantly affected (Supplementary figure 11B). Recent studies suggest potential cross-talk between siRNAs and piRNAs in insects, with maternally inherited siRNAs initiating piRNA cluster formation in *D. melanogaster* (Luo *et al*., 2023), and viral piRNAs appearing after viral siRNAs in *Ae. aegypti* (Ma *et al*., 2021). Our data, however, cannot confirm the role of siRNAs in piRNA biogenesis.

Since both endo-siRNAs and piRNAs are produced in the same tissues, there is likely a degree of redundancy between the two systems (Saito and Siomi, 2010). Indeed, both small RNA pathways have been shown to have redundant functions in general suppression of TE expression in *D. melanogaster* somatic tissues, where Piwi-mediated establishment of TE silencing during embryogenesis is largely sufficient to repress TE expression in the adult fly, and the siRNA pathway is capable of suppressing TEs in a *Piwi* knockout line (Beek *et al*., 2018). Anecdotally, the higher proportion of TEs present in the genomes of *Aedes* mosquitoes compared to *Drosophila* coincides with an expansion of *Piwi* genes, suggesting diversification and specialization of the piRNA pathway (Campbell *et al*., 2008; Lewis, Salmela and Obbard, 2016). It is possible that mosquitoes from this genetic background, which are viable and fertile despite a dysfunctional *Dcr2* gene (Merkling *et al*., 2023), are further less reliant on the endo-siRNA pathway for TE regulation. The lack of *Dcr2* in this genetic background also had little impact on vector competence for arboviruses, indicating a less significant role for siRNAs in antiviral defense than previously thought (Merkling *et al*., 2023). Nonetheless, the differential expression of individual TE families seen in the *Dcr2* mutant suggests varying degrees of sensitivity to TE silencing mechanisms, prompting future detailed investigation of individual families.

We previously observed that the *Dcr2* mutant line exhibits a range of modest fitness defects, such as slower development, increased pupal mortality, smaller adult body size, and reduced adult survival (Merkling *et al*., 2023). It is possible that some of these fitness defects reflect the modified patterns of TE expression, although additional investigation is required to support this hypothesis. Artificially stimulating retrotransposon activity was found to promote aging in *Drosophila* (Rigal *et al*., 2022). In other organisms, somatic TE mobilization is involved in various health defects such as cancer, aging and neurodegenerative diseases (Bourque *et al*., 2018).

In conclusion, our data are consistent with an auxiliary role of endo-siRNAs in silencing TE expression in *Ae. aegypti*, with individual TE families in different tissues displaying differing responses to a dysfunctional siRNA pathway in this genetic background. Unlike *D. melanogaster* (Chung *et al*., 2008; Czech *et al*., 2008; Ghildiyal *et al*., 2008), *Ae. aegypti* thus largely relies on other mechanisms than the siRNA pathway to control somatic TE expression.

## Acknowledgements

We thank Catherine Lallemand for assistance with mosquito rearing and Rachel Legendre for contribution to the transcriptomic analysis. We are grateful to Ronald van Rij and Rebecca Halbach for their advice about piRNA analyses. RNA-seq library preparation and sequencing was performed by the Biomics platform (C2RT, Institut Pasteur, Paris, France) supported by France Génomique (ANR-10-INBS-09) and IBISA. This work was funded by the French Government’s Investissement d’Avenir program, Laboratoire d’Excellence Integrative Biology of Emerging Infectious Diseases (grant ANR-10-LABX-62-IBEID to L.L.), and Agence Nationale de la Recherche (grant ANR-18-CE35-0003-01 to L.L.). A.B. was supported by a stipend from the Pasteur – Paris University (PPU) International PhD Program. The funders had no role in study design, data collection and analysis, decision to publish, or preparation of the manuscript.

## Author contributions

Conceived and designed the study: A.B., H.L.-M., J.D., M.-C.S., L.L.

Collected the data: A.B., H.B., L.F.

Contributed materials or analysis tools: A.B.C., S.H.M., M.C.-S.

Performed the analyses: A.B., H.L.-M., H.V., J.D., L.L.

Wrote the manuscript: A.B., L.L.

Acquired the funding: A.B., M.-C.S., L.L.

## Supplementary materials

**Supplementary figure 1.**
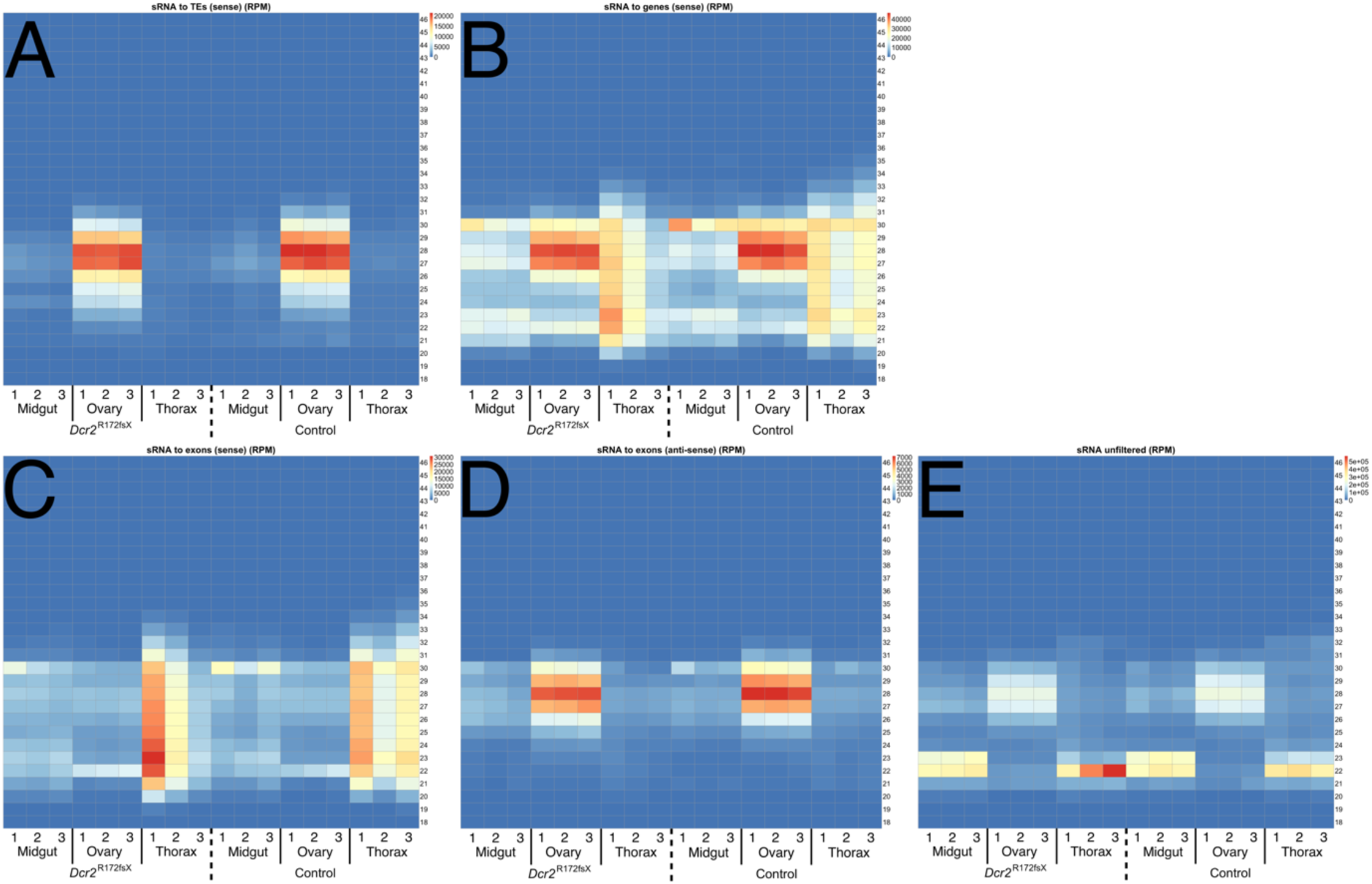
– Digital northern blots of small RNA-seq data. Size distributions of reads filtered for rRNA, miRNA, tRNA, snRNA, and snoRNA genes mapping to TEs on the sense strand (A), mapping to whole genes (exons, introns, and untranslated regions [UTRs]) on the sense strand (B), mapping to the sense strand of gene exons (C), mapping to the antisense strand of gene exons (D), and unfiltered reads (adapter-trimmed only) (E). Numbers underneath each lane represent the three biological replicates. Thorax samples in both the *Dcr2* mutant (lanes 7-9) and in the control mosquitoes (lanes 16-18) show an overabundance of reads of various sizes mapping to the sense strand of genes and, in particular, the sense strand of exons. Color corresponds to the RPM (non-adjusted by miRNA RPM).

**Supplementary figure 2.**
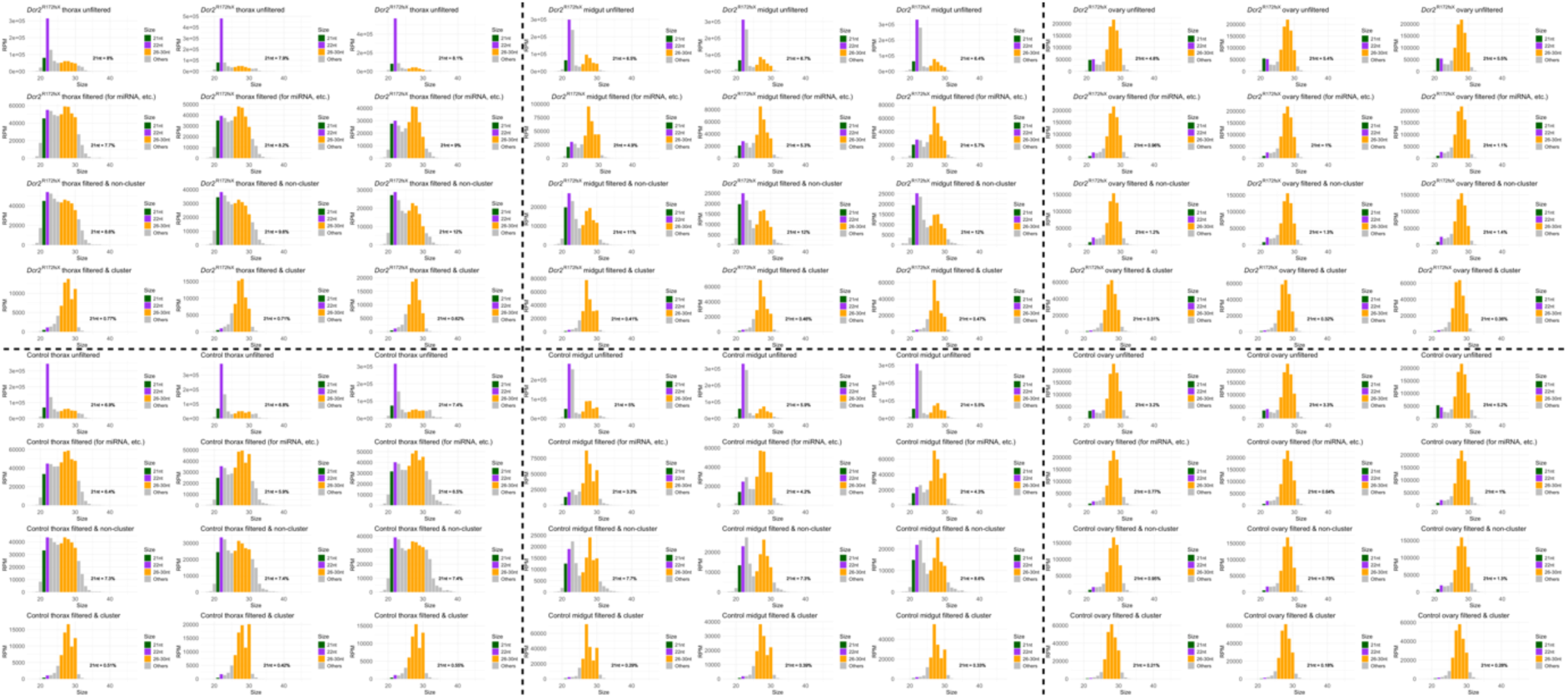
– Size distributions of reads in each sample following different levels of filtering. *Dcr2* mutant samples are shown in the top half of the figure, and control mosquito samples are shown in the bottom half. Thorax samples are on the left, midgut samples in the center, and ovary samples on the right. Within each partitioned region, the three columns of plots correspond to the three biological replicates. The rows correspond to reads subset based on filtering: First row – adapter-trimmed only (“unfiltered”), second row – filtered for reads mapping to miRNA, tRNA, snRNA, and snoRNA genes, third row – further filtered for reads mapping to annotated piRNA clusters, fourth row – only reads that map to piRNA clusters but not to any miRNA, tRNA, snRNA, or snoRNA genes. The percentage of 21nt reads is added to each plot. When filtered for small RNA genes, a ‘block’ of reads can be seen in thorax samples, attributable to RNA degradation, while a piRNA-sized ‘hump’ can be seen most clearly in ovary samples, but also in midgut samples. Unfiltered reads from somatic tissues display a clear domination of the library by miRNA-sized reads, which are filtered away following intersection with annotated small RNA genes.

**Supplementary figure 3.**
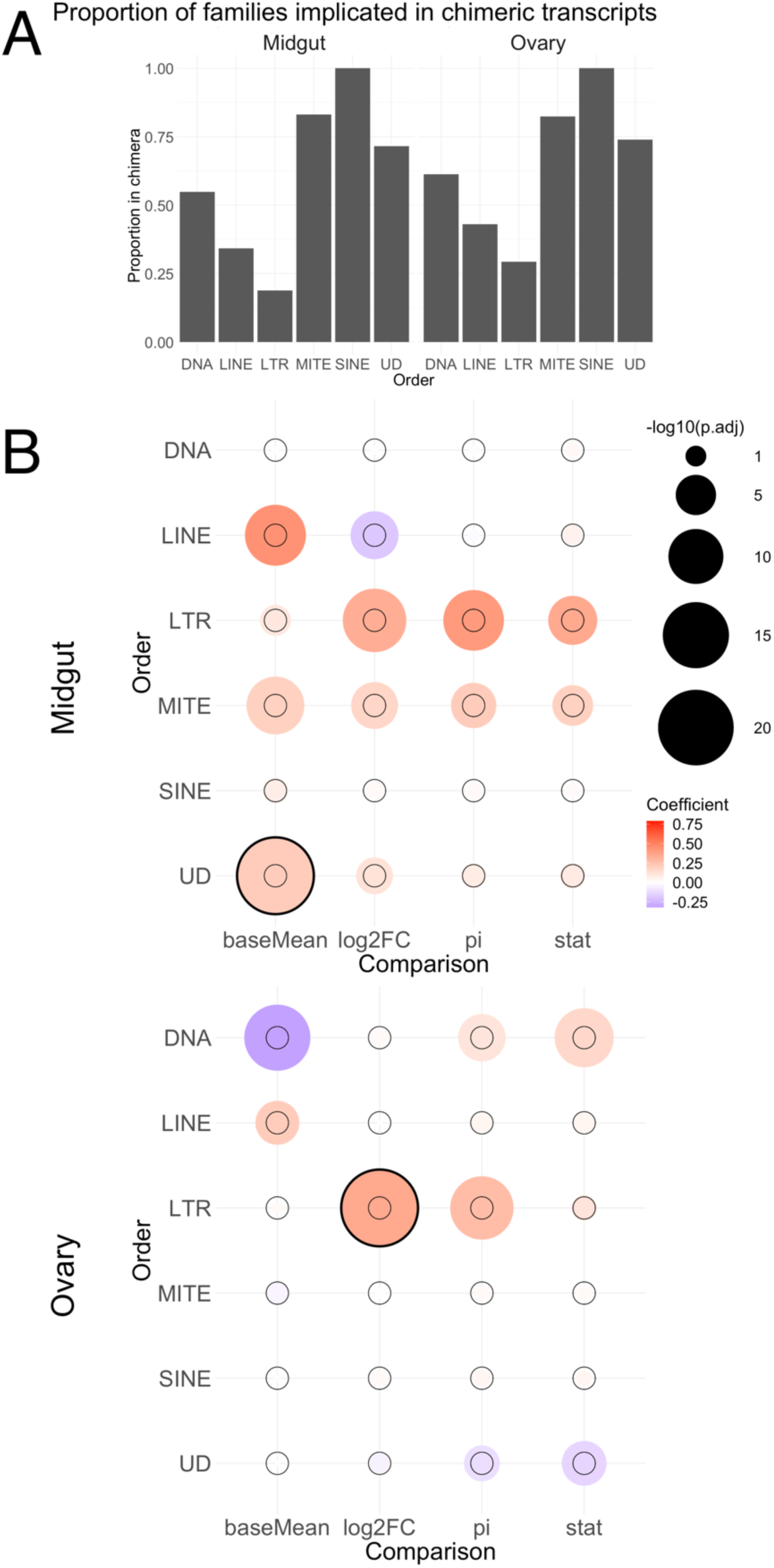
– MITEs and UDs are often found in chimeric transcripts. (A) Proportion of TE families in each order implicated (i.e., minimum 2 reads from all biological replicates combined supporting a given gene-TE association) in chimeric transcripts for midgut and ovary RNA-seq data. (B) Matrix of regression coefficients for midgut and ovary RNA-seq data. The parameters baseMean (number of counts normalized by median of ratios in control mosquitoes), the log_2_ fold-change (log2FC) in the *Dcr2* mutant, a combined metric (Xiao *et al*., 2014) for base expression and log_2_ fold-change (pi), and the DESeq2 statistic (stat) for genes implicated in chimeric transcripts were regressed as a function of the same parameter for implicated TEs (one regression for each organ, order, and parameter combination). A thin circle in the center of each grid intersection denotes the limit for statistical significance (p=0.05) of the slope coefficient for the regression. Negative-log_10_-transformed adjusted p-values above 20 are denoted with a thick outer circle. Significant positive coefficients for comparisons of all parameters are seen for LTR, MITE, and UD transposons in the midgut. Since most (>70%) of MITE and UD families are also implicated in gene-TE chimeric transcripts, some of the expression of these orders may be attributed to the expression of adjacent genes. For LTR transposons, positive regression coefficients are seen both in the midgut and the ovaries. However, the fraction of LTR transposons implicated in chimeric transcripts is much smaller (<20% for the midgut, <30% for the ovaries).

**Supplementary figure 4.**
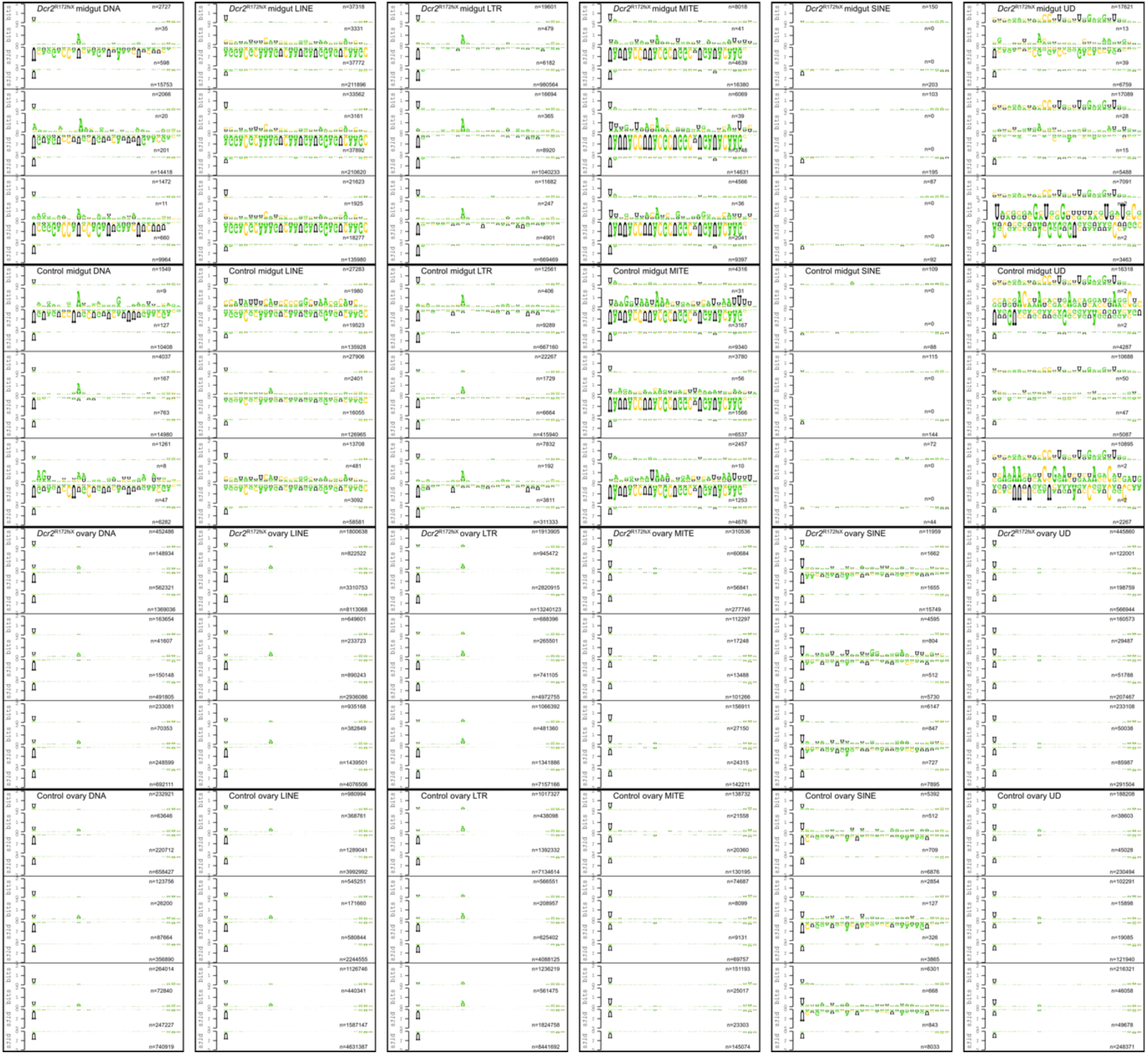
– Logo plots show 1U and 10A bias in putative piRNAs only for certain TE orders. Logo plots were constructed from 26-30nt reads mapping to the sense and antisense strands of TEs. Each column of logo plots in the figure corresponds to a TE order (DNA, LINE, LTR, MITE, SINE, UD=undetermined). The top half of the figure shows logo plots for the midgut samples, while the bottom half shows logo plots for the ovary samples. Within each half, the top half correspond to samples from the *Dcr2* mutant line, while the bottom corresponds to samples from control mosquitoes. Within each condition-organ-order partition, the three sets of four logo plots correspond to the three biological replicates. Each set is composed of four logo plots in the order top to bottom: 1) reads mapping to sense strand; 2) putative secondary piRNAs, i.e., reads mapping to sense strand and overlapping a putative primary piRNA in the secondary position (downstream of the corresponding putative primary piRNA) by 10 nt; 3) putative primary piRNAs engaged in the ‘ping-pong’ cycle, i.e., reads mapping to the antisense strand and overlapping a putative secondary piRNA in the primary position (upstream of the corresponding secondary piRNA) by 10 nt; 4) reads mapping to the antisense strand. The number of reads used to construct the logo is specified on the right side in each plot.

**Supplementary figure 5.**
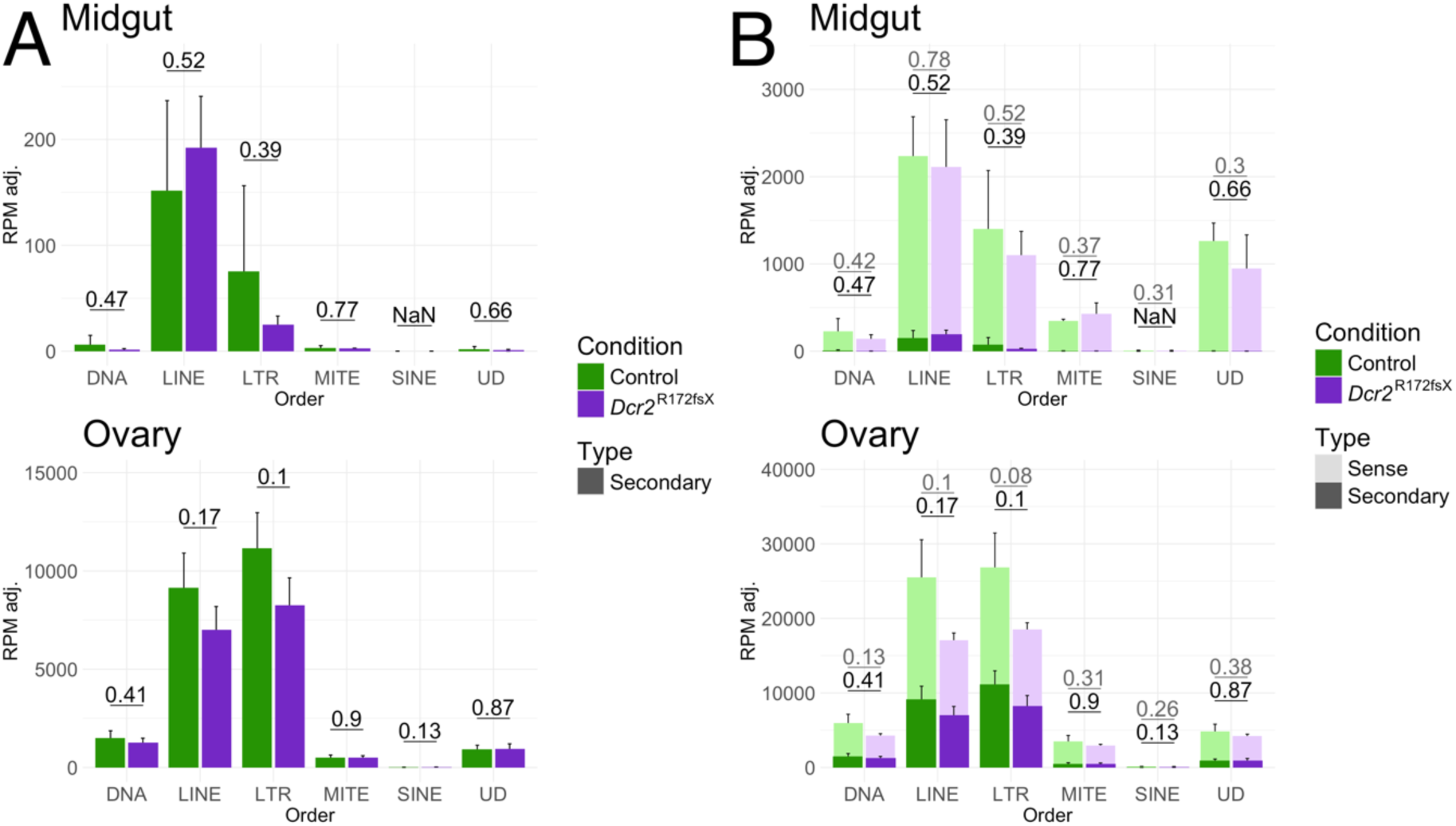
– Secondary piRNA abundance is equivalent between the *Dcr2* mutant and control lines. (A) Amounts of putative secondary piRNAs for each TE order for midguts (top) and ovaries (bottom). The numbers above the bar plots indicate p-values obtained with Welch’s t-test. For SINE, no secondary piRNAs were detected in the midgut samples. (B) Amounts of sense piRNA-sized reads for each TE order in midguts (top) and ovaries (bottom). The fraction of these that are considered putative secondary is highlighted in dark. Grey numbers above the bars indicate p-values obtained with Welch’s t-test for the amount of sense piRNA-sized reads. Black numbers indicate p-values for putative secondary piRNAs.

**Supplementary figure 6.**
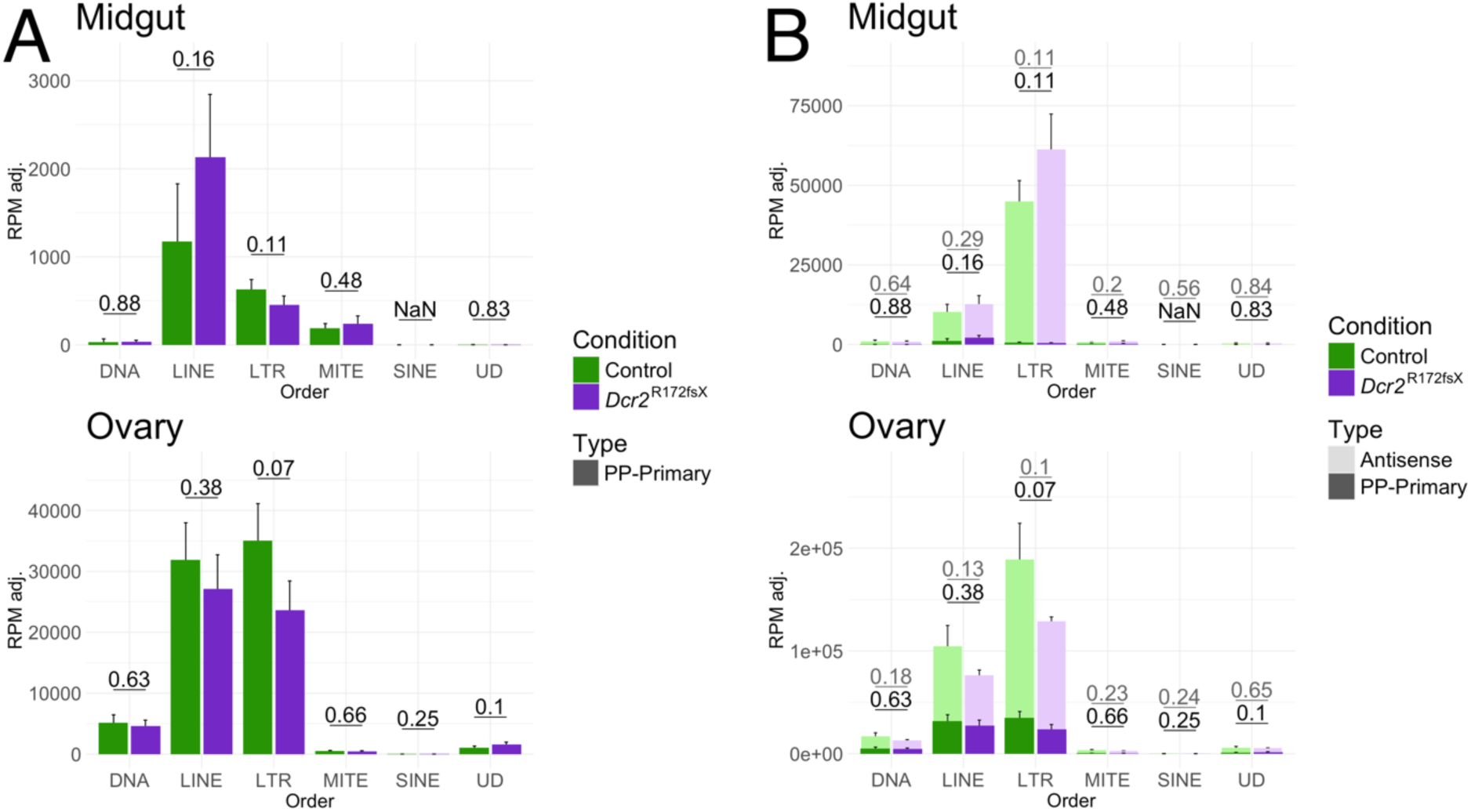
– Sense and ‘ping-pong’-interacting primary piRNA abundance is equivalent between the two mosquito lines. (A) Amounts of putative primary piRNAs identified by 10nt overlap-based analysis of ping-pong signature (PP-primary) for each TE order for midgut (top) and ovary (bottom) samples. The numbers above the bar plots indicate p-values obtained with Welch’s t-test. For SINE, no 10nt overlaps and thus no ‘ping-pong’-interacting primary piRNAs were detected in the midgut samples. (B) Amounts of antisense piRNA-sized reads for each TE order in midgut and ovary samples. The fraction of these that are considered putative primary piRNAs identified through 10nt overlap-based analysis of ping-pong signature (PP-primary) is highlighted in dark. Grey numbers above the bars indicate the p-value for the amount of antisense piRNA-sized reads. Black numbers indicate p-values for putative ‘ping-pong’-interacting primary piRNAs obtained with Welch’s t-test.

**Supplementary figure 7.**
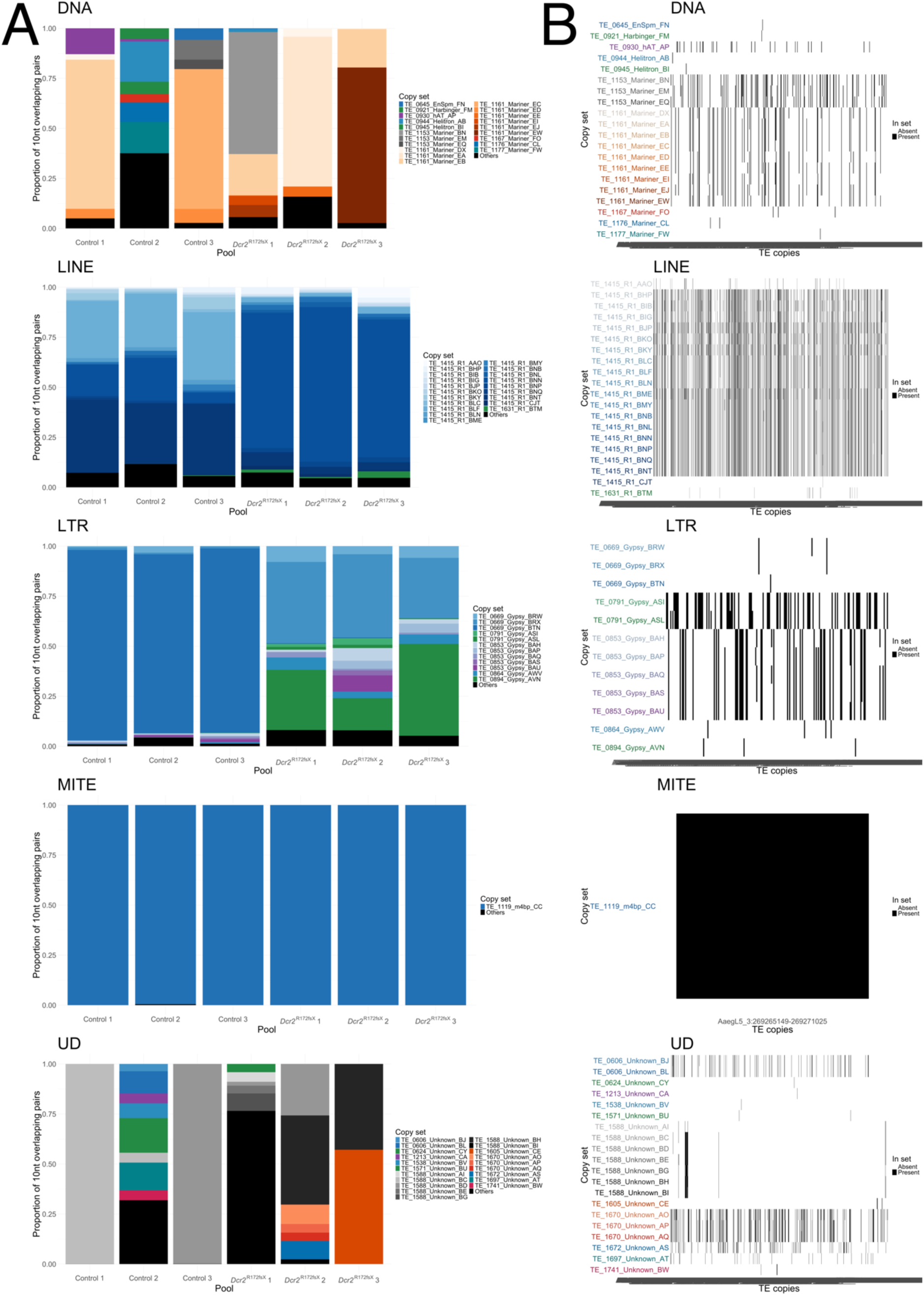
– Midgut piRNA-sized reads overlapping each other by 10 nt originating from LTR transposons are dominated by a single copy in wild-type mosquitoes. (A) The genomic origin of reads involved in 10nt overlaps in the midgut. For each TE order and replicate pool, the proportion of 10nt overlaps originating from a given set of TE copies. The letter index following the family name for each set of copies is arbitrary. For each order, up to 20 copy sets ordered by their contribution to total 10nt overlaps, or those necessary to reach 90% of total 10nt overlaps across all samples, were assigned colors. All other copy sets were assigned black and grouped into ‘Others’. (B) The copy composition of each set shown in (A). Individual copies are alphabetically ordered by genomic position on the x-axis.

**Supplementary figure 8.**
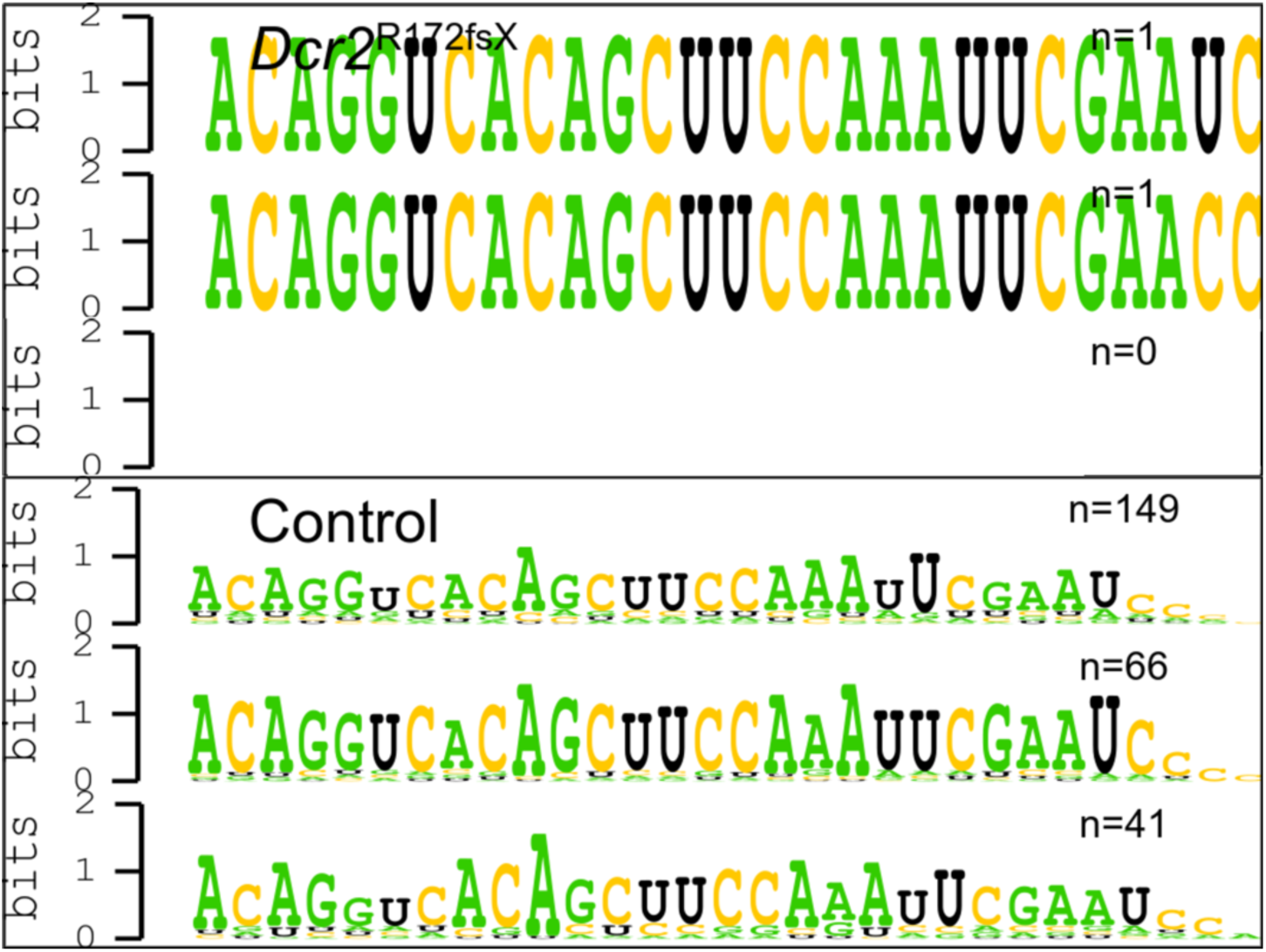
– Sequence composition of secondary piRNAs originating from the single locus mainly contributing to the 10nt overlaps among LTR transposons in the midgut of control mosquitoes. Logo plots derived from sequences mapping to the set of copies TE_0669_Gypsy_BTN, consisting of one single copy as seen in Supplementary figure 7. Top half of figure shows logo plots from the three replicates of the *Dcr2* mutant mosquitoes and bottom half shows the three replicates of the control mosquitoes.

**Supplementary figure 9.**
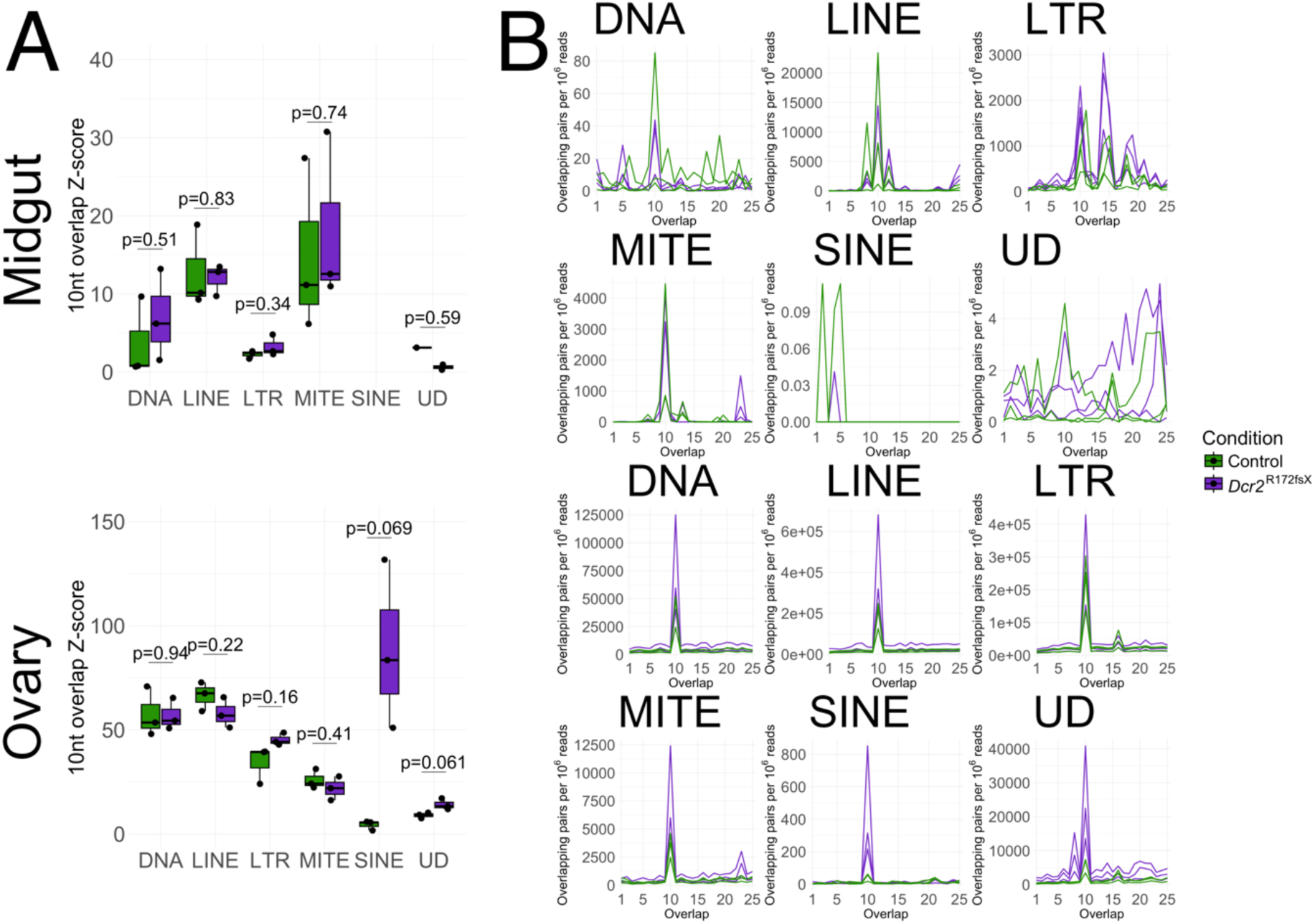
– Excluding the single locus contributing to the vast majority of 10nt overlaps for LTR transposons in the midgut abolishes the only difference between the *Dcr2* mutant and control lines. (A) Box plots of 10nt overlap Z-scores among 26-30nt sense and antisense reads mapping to TEs in midguts (top) and ovaries (bottom). P-values indicated above the box pairs were obtained using Welch’s t-test. (B) Frequency of overlaps among sense and antisense reads by a given number of nt for midguts (top two rows) and ovaries (bottom two rows). Reads mapping to the copy of TE_0669_Gypsy that dominated the overlaps in control mosquito midguts were excluded from the plot for LTR transposons for the midgut samples.

**Supplementary figure 10.**
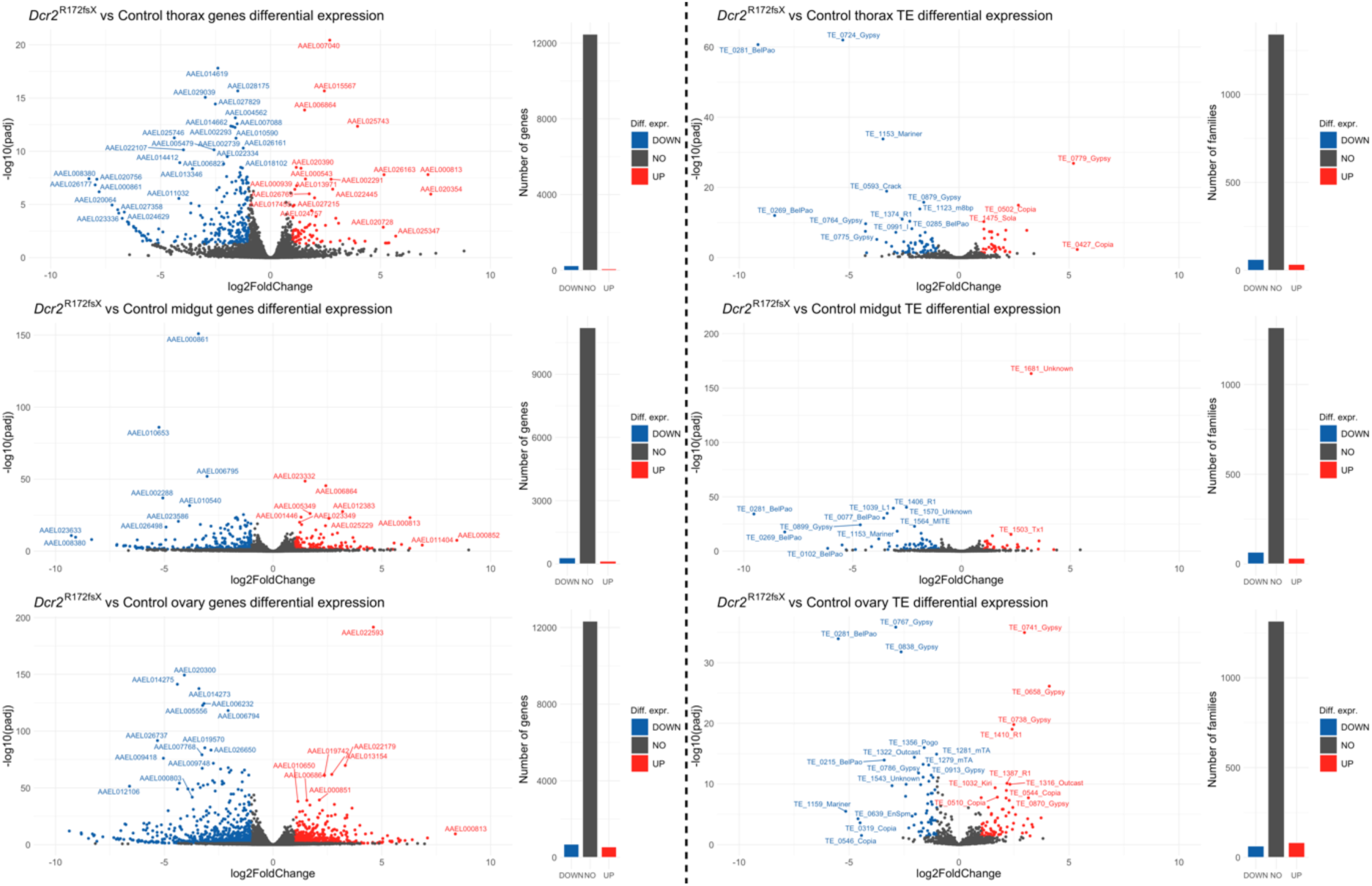
– Volcano plots and summary bar plots for differentially expressed genes and TE families show a wide perturbation of expression. Plots to the left of the partition correspond to genes, while plots to the right of the partition correspond to TEs. The three rows of plots correspond to the organs (ordered top to bottom): thorax, midgut, and ovary. The bar plots to the right of the volcano plot summarize the number of depleted (DOWN), non-differentially expressed (NON), and enriched (UP) genes or TEs in the *Dcr2* mutant line relative to the control line. Genes or TE families with an absolute log_2_ fold-change >1 and an adjusted p-value <0.05 are colored according to the direction of their differential expression (red: enriched; blue: depleted; grey: not differentially expressed).

**Supplementary figure 11.**
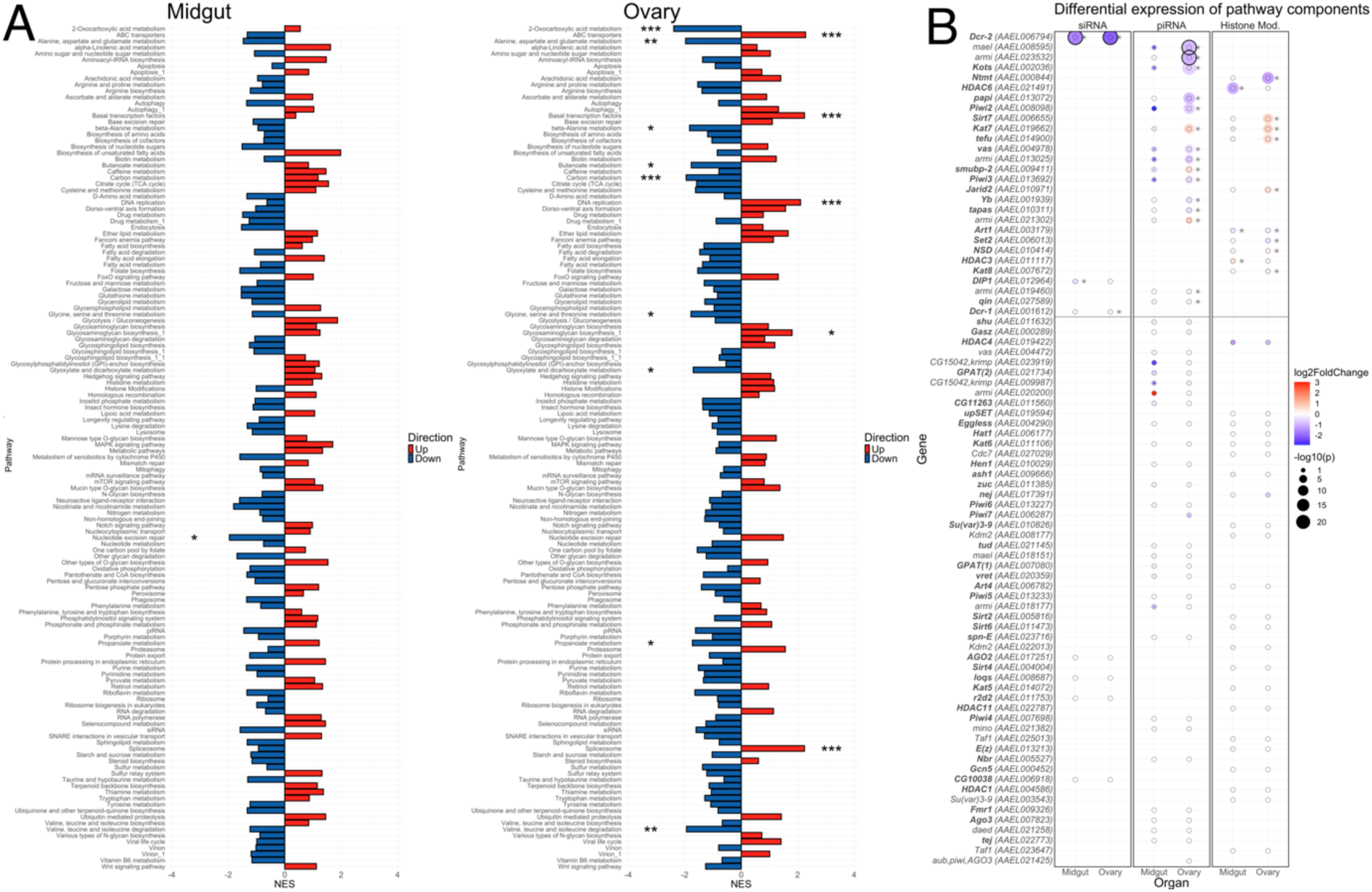
– Gene set enrichment analysis for genes shows multiple differentially regulated pathways in the ovaries. (A) GSEA for annotated KEGG pathways as well as manually added pathways ‘siRNA’, ‘piRNA’, and ‘Histone Modification’ for midgut (left) and ovary (right) samples. Significant enrichments or depletions are labelled with asterisks indicating the false discovery rate (*<0.05, **<0.01, ***<0.001). (B) Differential expression of *Ae. aegypti* homologs of *D. melanogaster* genes associated with the siRNA-, piRNA-, and histone modification-related pathways. Gene names in bold font indicate either a unique homolog, or a homolog with an annotated homologous function. Within each colored circle, a thin black circle indicates the threshold for statistical significance (p=0.05). The size of the colored circle corresponds to the significance level for differential expression expressed as –log_10_(p-value). Significantly differentially expressed genes are highlighted with an asterisk in the plot. Genes with a negative-log_10_-transformed p-value greater than 20 have their colored circles surrounded by a thick outer circle. A horizontal line separates the genes with any significant differential expression from those with none.

**Supplementary figure 12.**
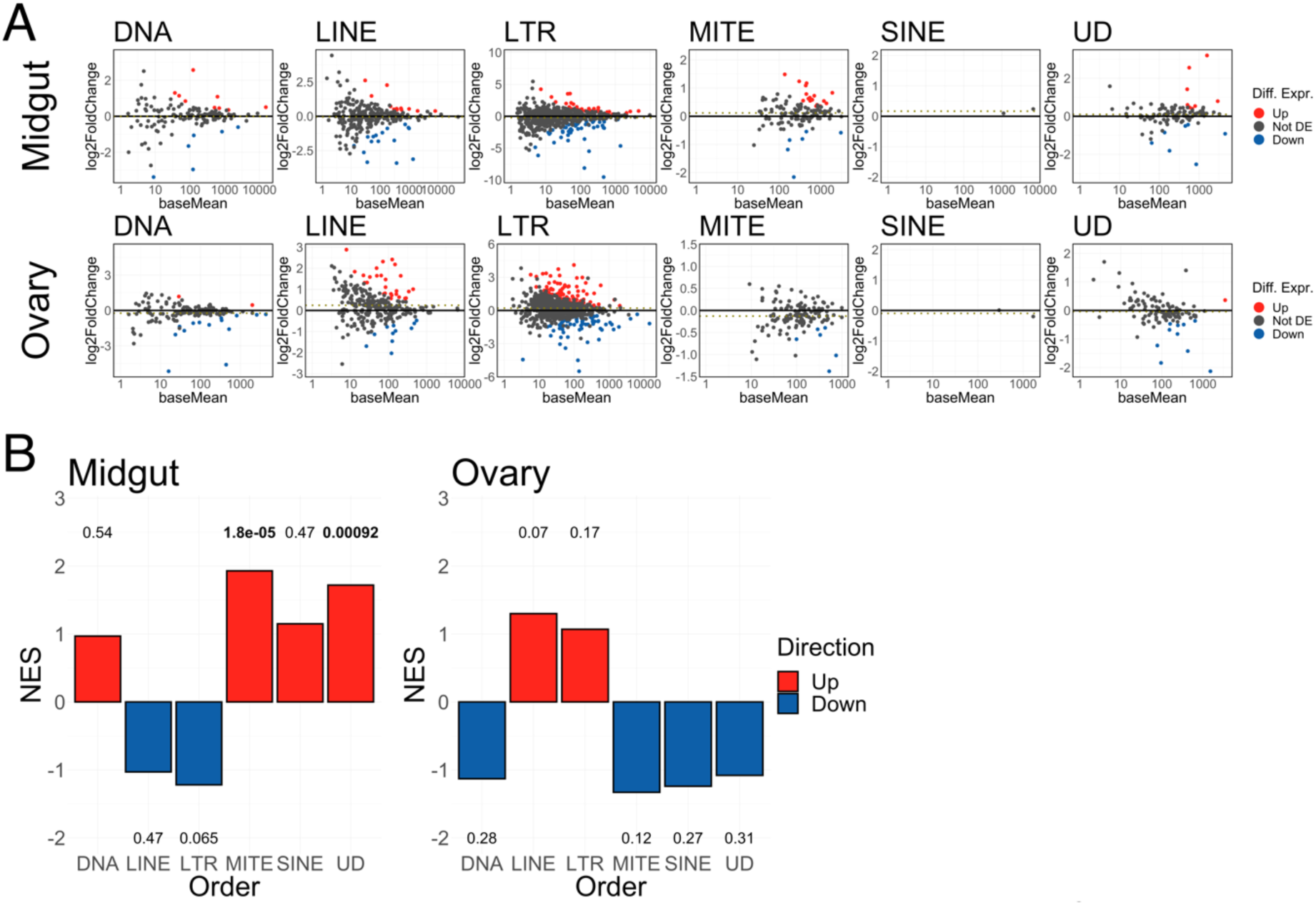
– TE expression for all orders (autonomous and non-autonomous). (A) MA plots of individual TE families grouped by order for midguts (top row) and ovaries (bottom row). The x-axis shows mean read counts normalized by the median of ratios and the y-axis shows log_2_ fold-change in the *Dcr2* mutant line. Families are colored according to their differential expression (red: enriched in mutant line; blue: depleted in mutant line; grey: not differentially expressed). The dotted line in the center of each plot represents the mean log_2_ fold-change. (B) Gene set enrichment analysis results. The height of each bar represents the normalized enrichment score (NES), i.e., the relative enrichment of the TE order compared to a random group of genes with the same size. Numbers above or below the bars indicate the false discovery rate for the enrichment (red bars) or depletion (blue bars) in the *Dcr2* mutant relative to the wild-type control.

**Supplementary figure 13.**
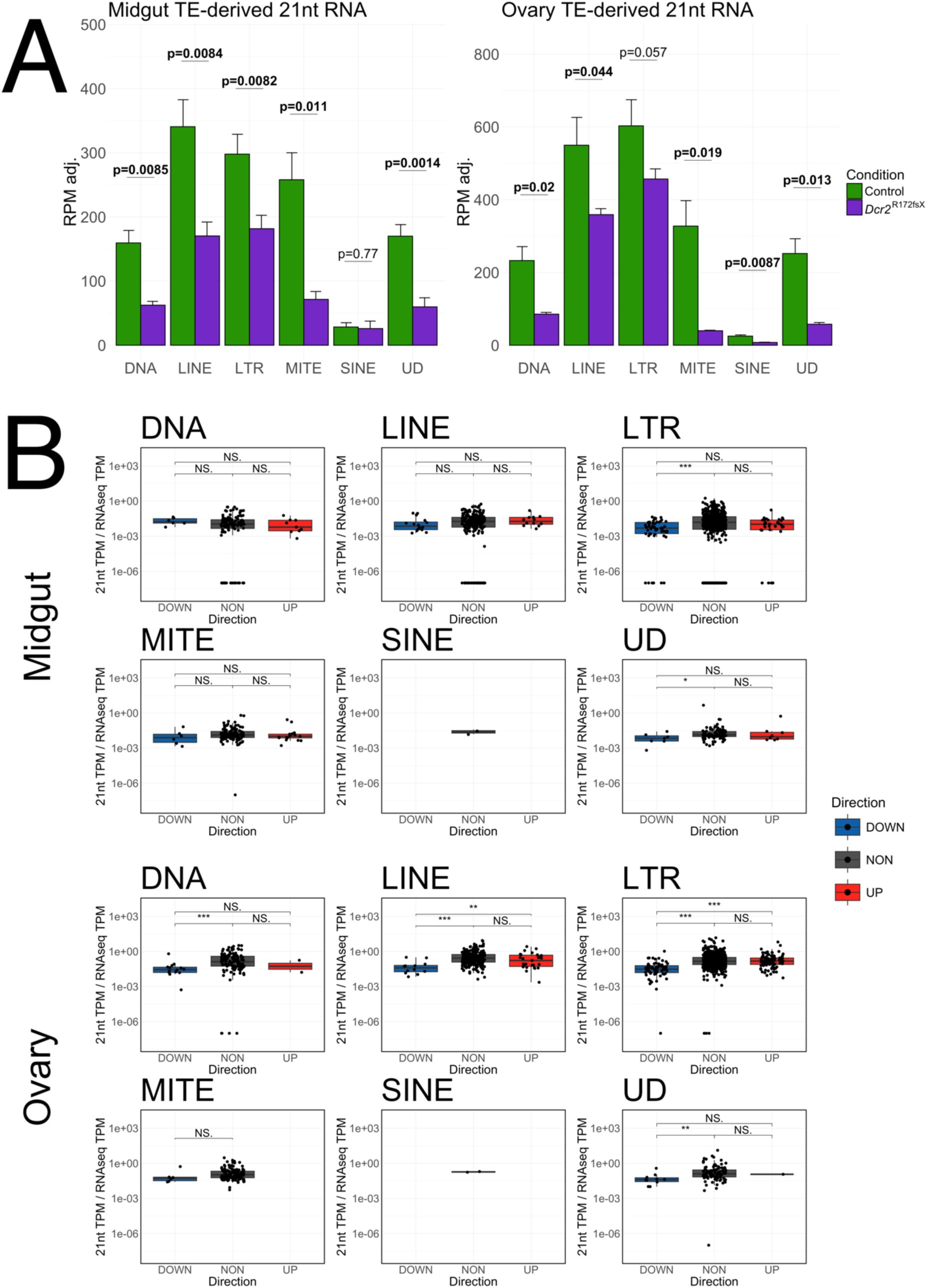
– Correlation between TE-derived 21nt RNAs and differential TE expression for all orders. (A) miRNA-adjusted reads per million (RPM) mapping to the different TE orders in the midgut (left plot) and ovaries (right plot) of *Dcr2* mutant and control mosquitoes. Error bars denote one standard deviation. Statistical significance was determined using Welch’s t-test (*p<0.05, **p<0.01, NS = non-significant). (B) Ratios between miRNA-adjusted 21nt RNAs expressed in transcripts per million (TPM) and RNA-seq TPM in the control mosquitoes for individual TE families depleted (Down), non-differentially expressed (Not DE), and enriched (Up) in the *Dcr2* mutant within each TE order for midguts (top two rows) and ovaries (bottom two rows). Statistical significance was determined using Wilcoxon rank sum test (*p<0.05, **p<0.01, ***p <0.001, NS = non-significant). Families with no detected transcripts were excluded. Families with detected transcripts but no detected 21nt RNAs are shown below the dashed line.

